# GM-CSF production by immune cells in steady state and autoimmune neuroinflammation mapped using fate reporting mice

**DOI:** 10.1101/2025.04.23.650273

**Authors:** Gholamreza Azizi, Javad Rasouli, Hamed Naziri, Michael V. Gonzalez, James Garifallou, Guang-Xian Zhang, Bogoljub Ciric, Abdolmohamad Rostami

**Author notes:** **Corresponding author:** A. Rostami, M.D. Ph.D. Department of Neurology, Thomas Jefferson University, 900 Walnut Street, Suite 300, Philadelphia, PA, 19107.

## Abstract

The full GM-CSF expression spectrum in immune cells remains unclear, while CD4□ T cells are the primary source. Using novel GM-CSF reporter/fate reporter transgenic mice, we tracked ongoing and past (YFP^+^) GM-CSF expression in various immune cells. GM-CSF was produced by diverse immune cells, including CD4^+^, CD8^+^, γδ T, NK, B, and CD11b^+^ cells, with expression patterns varying by cell type and organ with liver CD4^+^ T cells and NK cells showing the highest expression history in both naïve and mice with EAE. GM-CSF expression was transient and permanently lost in most cells over time. In a mouse model of multiple sclerosis, effector memory CD4□ T cells were the dominant CNS GM-CSF source, with higher expression than in other organs. CD4^+^YFP^+^ T cells, strongly expressing CXCR6, produced multiple cytokines. Transcriptomic analysis showed distinct gene expression profiles in effector memory CD4^+^ T cells compared to naïve cells. YFP□ Tregs represent functionally distinct subsets mirroring effector Th cells, expressing cytokines associated with Th lineages, especially during neuroinflammation. These findings identified distinct GM-CSF cellular sources across organs, highlighting a transient tissue microenvironment influence on GM-CSF production linked to CXCR6 expression.

## Introduction

Granulocyte macrophage-colony stimulating factor (GM-CSF, also known as CSF2) is a hematopoietic growth factor and immune modulator that promotes the proliferation and differentiation of bone marrow progenitor cells into myeloid lineages (1, 2). GM-CSF’s heteromeric cell-surface receptor, comprising GM-CSFRα and GM-CSFRβc subunits, mediates its biological effects and is expressed on multiple cell types, including monocytes, macrophages, neutrophils, endothelial cells, and alveolar epithelial cells (3, 4, 5). While GM-CSF is typically expressed at low levels in healthy tissues (6), its production increases during inflammation and autoimmune diseases (5, 7, 8, 9).

GM-CSF is primarily produced by lymphoid and myeloid cells, particularly in response to inflammatory stimuli (8, 10). Among these cell types, T cells are recognized as likely the most relevant cellular source of GM-CSF in the pathogenesis of multiple sclerosis (MS) (11, 12). Although the contribution of GM-CSF to the encephalitogenicity of CD8^+^ T cells remains unknown (12), CD4^+^ T cells are viewed as the predominant source of GM-CSF in neuroinflammation (8). Initially, Th1 and Th17 cells were thought to be the major producers of GM-CSF, playing a crucial role in the pathogenesis of experimental autoimmune encephalomyelitis (EAE) (11, 13, 14) in the animal model of MS. However, recent studies identified a distinct GM-CSF-producing Th subset, ThGM, that is probably involved in EAE development (8, 10, 15). MS patients exhibit an expansion of GM-CSF-secreting CD4^+^ T cells, indicating their pathogenic role in neuroinflammation (12, 16). Several non-T cell types also produce GM-CSF, contributing to inflammatory responses. These include B cells, innate lymphoid cells (ILCs), and resident tissue cells such as endothelial cells, fibroblasts, and epithelial cells (16, 17, 18). Collectively, these findings underscore the diverse cellular sources of GM-CSF and their roles in shaping inflammatory environments across different disease contexts.

GM-CSF expression in vivo has been partially explored (10), but the current study substantially deepens current knowledge using a fate reporter system, which can determine correlations between prior and ongoing GM-CSF expression. We identified distinct expression patterns across various organs by examining the localization and fate of GM-CSF-expressing cells in vivo. We characterized GM-CSF expression in both steady state and neuroinflammation, focusing on its production by CD4□ T cell subsets and other tissue-resident immune cells. Our findings highlight effector memory CD4□ T cells as the primary source of GM-CSF during neuroinflammation. Additionally, we observed that GM-CSF-expressing T cells co-express CXCR6, suggesting a potential link between GM-CSF production and tissue residency. Notably, there is a strong correlation between the history of GM-CSF expression by CD4^+^ T cells and its overall phenotype. Specifically, effector memory CD4^+^ T cells that have previously expressed GM-CSF and those that never did have vastly different transcriptomes. This indicates that factors driving/regulating GM-CSF expression by CD4^+^ T cells have additional widespread effects. Furthermore, our results show that GM-CSF expression in CD4□ T cells is transient and permanently lost in most of the cells that previously expressed GM-CSF.

## Results

### Varying levels of past and current GM-CSF expression in immune cells across different organs

A wide range of immune cells, including T, B, innate lymphoid, and myeloid cells, can express GM-CSF (19). However, the numbers and phenotypic properties of GM-CSF-producing immune cells in various tissues and their fates in steady-state and inflammation remain incompletely understood. To address this knowledge gap, we used our *Gr/fr* transgenic mice to investigate GM-CSF expression across different immune cell populations. Multiple immune cell types produced GM-CSF, including CD4^+^, CD8^+^, γδ T, NK, B, and CD11b^+^ cells (**Fig. 1A**; general gating strategy is shown in **Fig. S1B**). Notably, there were substantial differences in past and current GM-CSF expression among immune cell types within the same anatomical location, as well as between the same cell types across different organs. For instance, while only a small proportion of B cells expressed GM-CSF, up to 80% of γδ T cells expressed it. Similarly, past GM-CSF production by CD8^+^ T cells ranged from a small percentage in the LNs to 70% in the SI (**Fig. 1B**).

**Figure 1.**
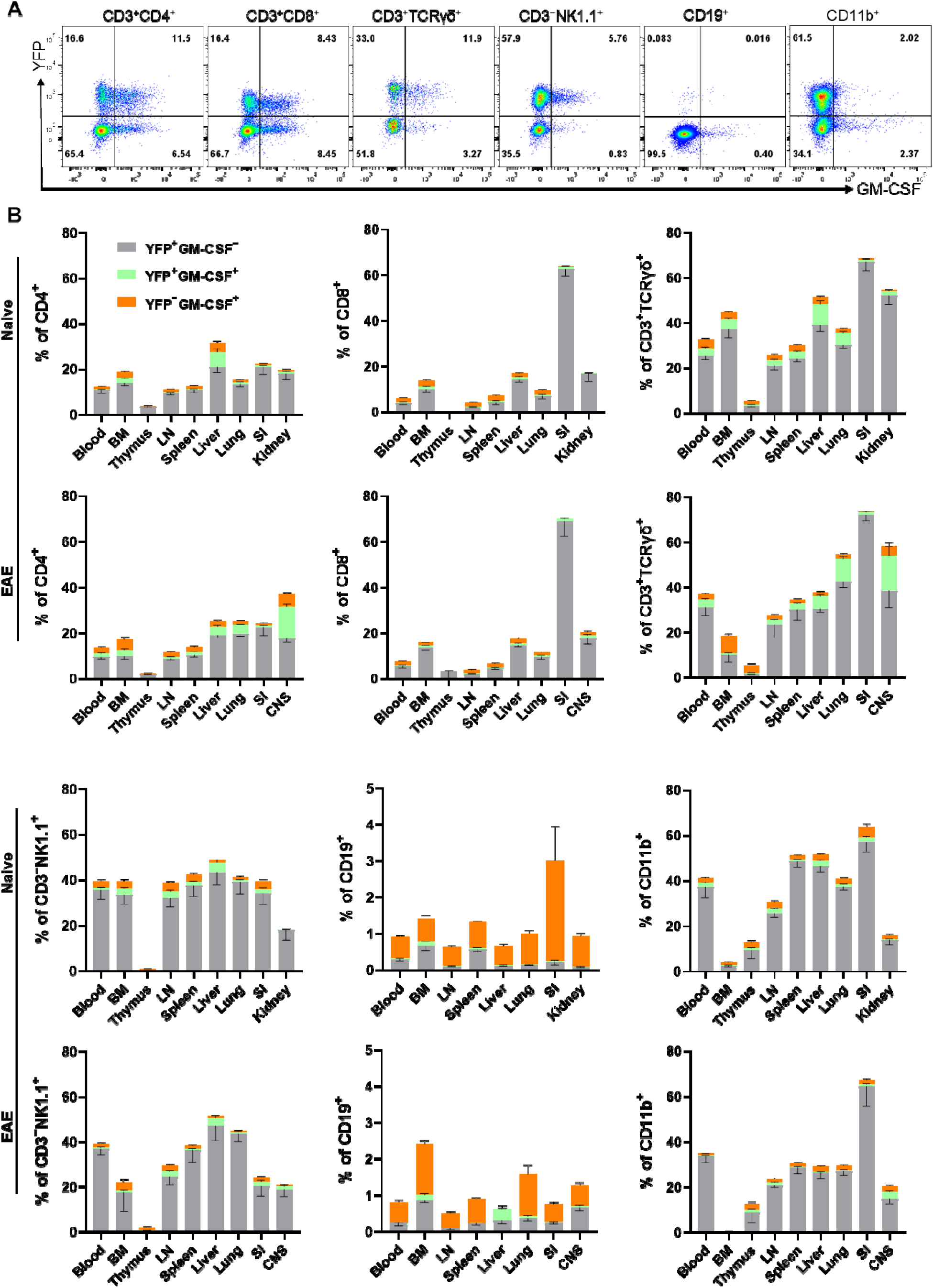
GM-CSF and YFP expression by immune cells from mouse organs. Mononuclear cells were isolated from the blood, BM, thymus, LNs, spleen, liver, lung, kidney (of naïve mice), SI, and CNS (of mice with EAE at the peak of clinical disease severity) from 2–3-month-old male and female Gr/fr mice. Cells were stimulated with PMA, Ionomycin, and GolgiPlug, and stained for CD45, lineage-specific surface markers, and GM-CSF. Cells were then analyzed by flow cytometry. YFP and GM-CSF expression was assessed within gated live CD45^hi^ populations. **(A)** Flow cytometry plots show YFP and GM-CSF expression in liver immune cells of naïve mice; a similar gating strategy was used for immune cells from other organs. **(B)** The stacked bar charts show the frequencies of YFP^+^ and GM-CSF^+^ cells among major immune cell types of naïve mice and mice with EAE. Data represents three independent experiments in naïve mice (total n = 13) and two independent experiments in mice with EAE (total n = 8). Error bars indicate the mean ± SEM. Abbreviations: LN- lymph nodes, BM - bone marrow, SI – small intestine, CNS- central nervous system.

In naïve mice, CD4^+^ T cells and NK cells in the liver exhibited higher levels of ongoing and past GM-CSF expression than in other organs, while CD8^+^ and γδ T cells in the SI displayed elevated GM-CSF expression. Additionally, CD11b^+^ cells in the spleen and liver showed greater GM-CSF expression than in other organs (**Fig. 1B**). The pattern of GM-CSF expression in mice with EAE, a model of CNS inflammation, was largely similar to that in naïve mice, except for reduced expression in CD11b^+^ cells. Notably, less than 1% of microglia displayed GM-CSF expression, either ongoing or in the past (**Fig. S1C**). In the CNS of EAE mice, approximately 40% of CD4^+^ T cells and 60% of γδ T cells expressed GM-CSF, with ongoing expression being notably higher compared to other organs (**Fig. 1B**). This is not surprising, as EAE involves infiltration of the CNS by highly activated immune cells.

The primary populations expressing YFP, indicative of prior GM-CSF production, include CD4^+^, CD8^+^, γδ T cells, and CD11b^+^ cells. Among CD45^hi^YFP^+^ cells, the dominant populations vary by organ. In the blood, spleen, and lungs of naïve mice, CD11b^+^ cells constitute the main YFP^+^ population, followed by CD4^+^ T cells. In the LNs and thymus, CD4^+^ T cells predominate (**Fig. 2A**). In the liver and kidney, both CD11b^+^ and CD4^+^ T cells represent the largest YFP^+^ populations. In the SI, γδ T cells and CD8^+^ T cells are the primary YFP^+^ populations. Within the CD45^hi^YFP^+^ population, NKT and B cells represent rare subsets, while NK cells account for 1– 8% across various organs. Additionally, a subset of CD3^+^γδ^−^ cells that did not express CD4 or CD8 was observed in different organs, mostly in the BM, kidney, liver, and lung (**Fig. 2**).

**Figure 2.**
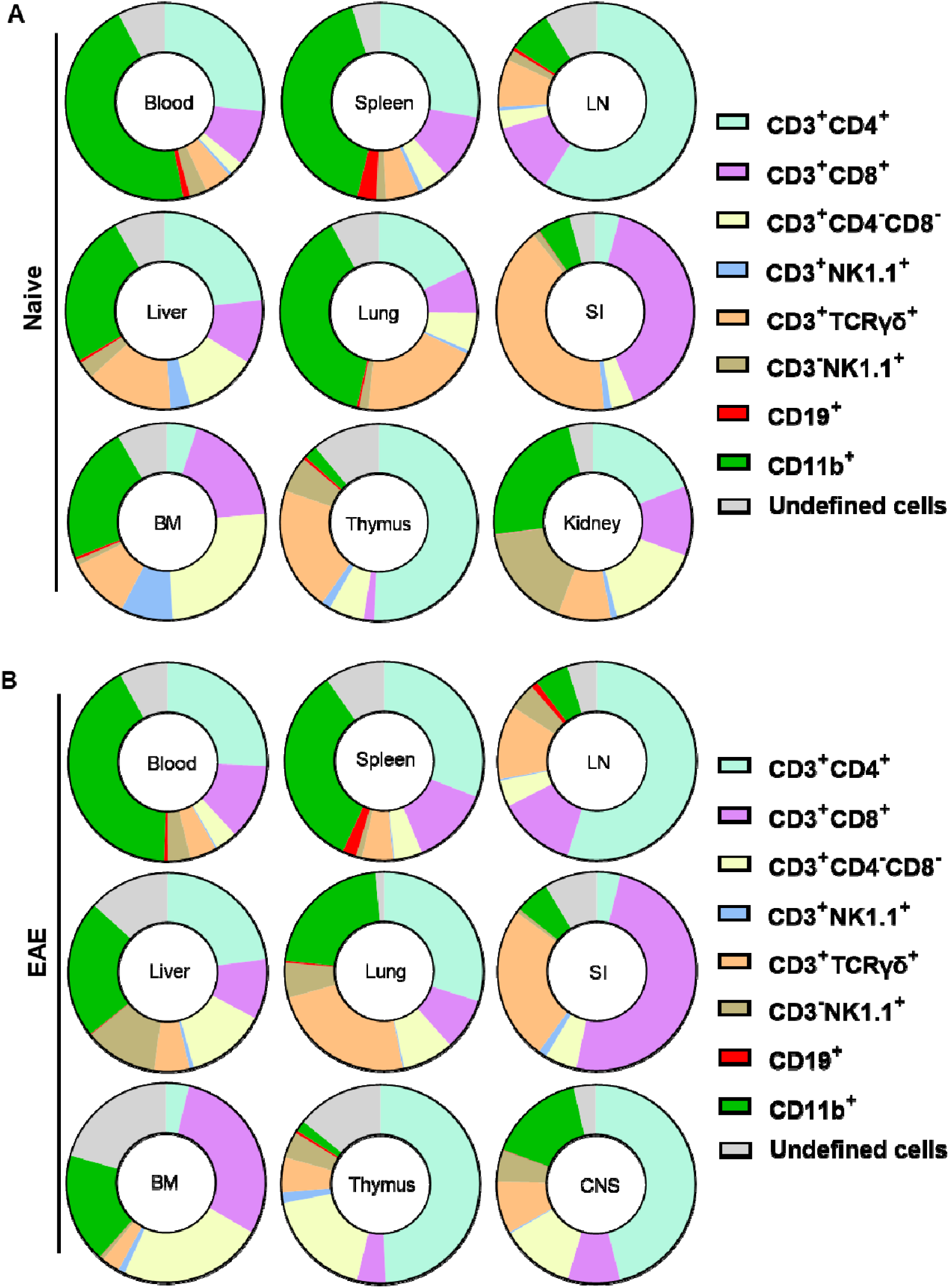
Proportions of cell types among CD45^hi^YFP^+^ immune cells of naïve (A) and (B) Gr/fr mice with EAE. Mononuclear cells were isolated from the blood, BM, thymus, LN, spleen, liver, lung, kidney (of naïve mice), SI, and CNS (of EAE mice at the peak of clinical disease) from 2–3-month-old male and female Gr/fr mice. Cells were stained for CD45, lineage-specific surface markers, and analyzed by flow cytometry. Proportions of YFP^+^ cells were determined within gated live CD45^hi^ populations. The pie charts show the frequencies of major cell types among the CD45^hi^YFP^+^ immune cells. Data is presented as the mean percentage from three independent experiments in naïve (total n = 13) and two independent experiments in EAE mice (total n = 8). The "undefined" cells include rare populations such as CD3□CD19□, CD3□CD19^low^, CD3□CD8^low^, CD3□CD4^low^, and CD3□CD4□CD8□ cells. Abbreviations: LN- lymph nodes, BM - bone marrow, SI – small intestine, CNS- central nervous system.

In EAE mice, the overall pattern of YFP expression remained similar to that of naïve mice, except in the lungs, where an increase in CD4^+^YFP^+^ cells rendered them the predominant population. In the CNS of EAE mice, CD4^+^ T cells were the most abundant YFP^+^ population (**Fig. 2B**).

Among CD11b^+^ cells, most YFP^+^ cells were macrophages (CD11b^+^CD11c^+^Ly6G^−^Ly6C^−^MHC-II^−^; **Fig. S2A**) in both naïve and EAE mice. Monocytes (CD11b^+^CD11c^−^Ly6C^hi^Ly6G^−^MHC-II^−^) and cDCs (CD11b^+^CD11c^+^Ly6G^−^Ly6C^−^MHC-II^+^) constituted minor CD11b^+^YFP^+^ populations. Notably, active GM-CSF expression was restricted to cDC (**Fig. S2B**). In both naïve and EAE mice, CD45^−^ cells did not express YFP, except for rare populations (<1%) in the lung and SI (**Fig. S2C**). In conclusion, diverse immune cells produce GM-CSF, with the primary sources being lymphoid cells, particularly CD4^+^ T cells, and macrophages among myeloid cells.

A particularly striking observation was that the vast majority of YFP^+^ cells of all types failed to re-express GM-CSF upon stimulation with PMA and Ionomycin. This suggests that GM-CSF expression is transient and that cells that used to express GM-CSF cease the expression irreversibly. Consequently, only a small subset of any immune cell type could express GM-CSF at any given time.

### Effector/memory CD4^+^ T cells are the main GM-CSF source in the lung and CNS during neuroinflammation

Given that T cells, especially CD4^+^ T cells were the major producers of GM-CSF, we focused our studies on these cells. In the organs of both naïve and mice with EAE, the frequencies of CD4^+^ T cells were higher than those of CD8^+^ T cells, except in the SI, where CD8^+^ T cells predominated (**Figs. S3A and B**). In lymphoid organs, blood, and lungs of naïve mice, conventional (Foxp3^−^) CD4^+^ and CD8^+^ T cell subsets were predominantly CD62L^+^CD44^−^, indicative of a naïve phenotype. In contrast, in the SI, the frequency of naïve cells was less than 5%. The proportion of CD62L^−^CD44^+^ cells, indicative of effector memory T cells, was higher in the liver, SI, and CNS. However, in the blood and lymphoid organs, only a small percentage (approximately 1-10%) of conventional CD4^+^ or CD8^+^ T cells were CD62L^−^CD44^+^. A shift toward CD62L^−^CD44^+^ cells was observed in all organs of EAE mice, most notably in the case of CD4^+^ cells in the CNS and lung (**Figs. S3C and D**), along with an increased frequency of MOG_35-55_-specific CD4^+^CD44^+^ T cells in the lung at 8 d.p.i. (**Figs. S3E and F**).

Using *Gr/fr* mice, we next investigated GM-CSF expression in subpopulations of conventional CD4^+^ and CD8^+^ T cells (**Fig. 3A**) across various organs of both naïve and EAE-affected mice. The highest frequencies of CD4^+^YFP^+^ cells were observed in the liver and CNS of naïve mice, with a noted increase in the lung and CNS following EAE immunization (**Fig. 3B**). For CD8^+^YFP^+^ T cells, the highest frequency was found in the SI, with no significant changes post-immunization. Within conventional CD4^+^ and CD8^+^ T cell subsets, YFP^+^ cells were most prevalent in the effector memory (CD62L^−^CD44^+^) subsets of both naïve (**Fig. 3C**) and EAE-affected mice (**Fig. 3D**), with higher frequencies in the lung and CNS of mice with EAE. There was little difference in proportions of CD8^+^YFP^+^ cells between naïve and EAE-affected mice. In conclusion, GM-CSF-expressing effector/memory CD4^+^ T cells were notably enriched in the lung and CNS of mice with EAE compared to naïve mice.

**Figure 3.**
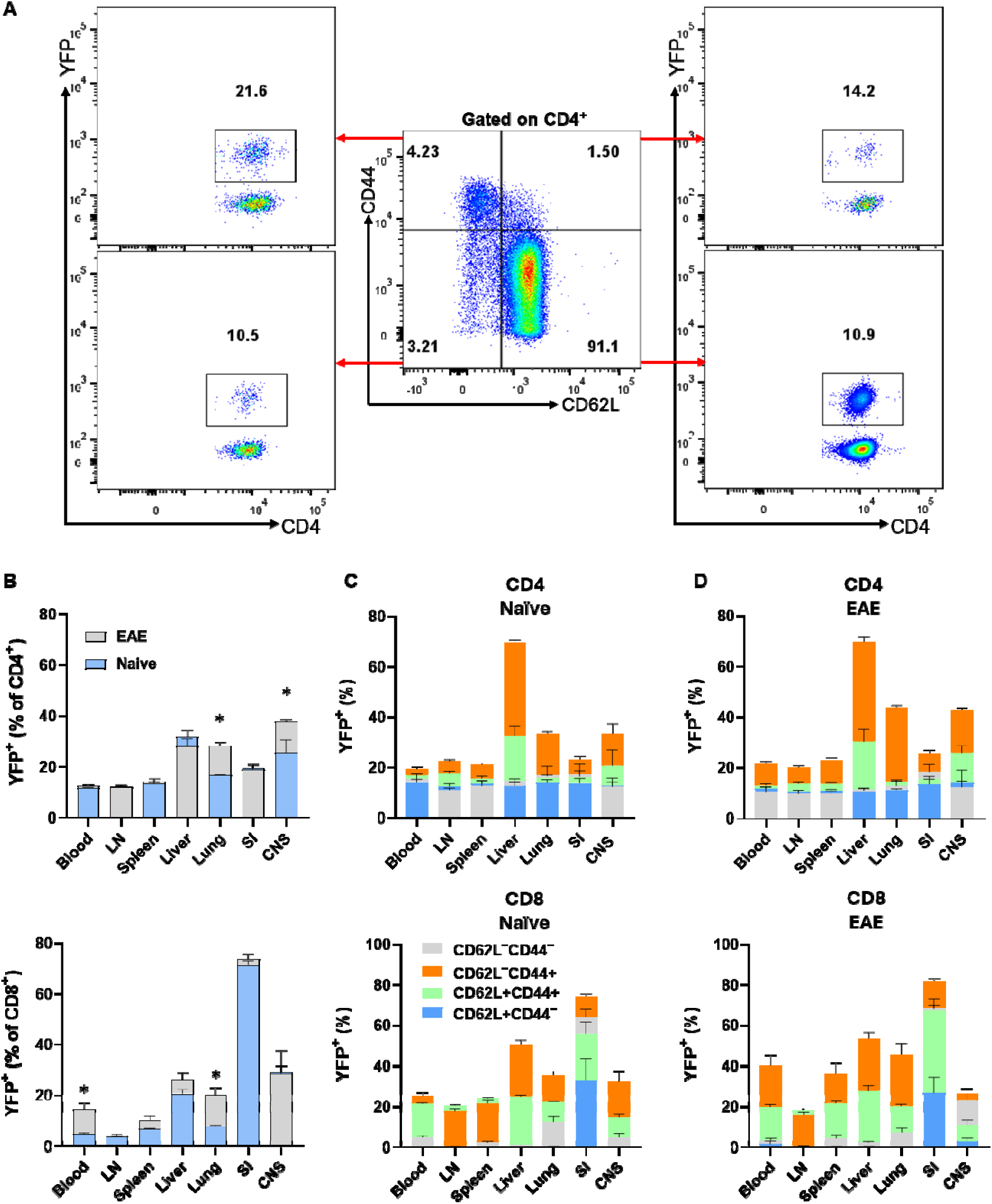
YFP expression by CD4^+^ and CD8^+^ T cells from various mouse organs. Gr/fr naïve and mice wit EAE (both male and female, aged 2–3 months) were sacrificed. Cells isolated from the blood, LN, spleen, liver, lung, SI, and CNS were stained and analyzed by flow cytometry for CD45, CD4, CD8, CD62L, CD44, and YFP expression. **(A)** The gating strategy for YFP expression analysis among CD62L^−^CD44^−^, CD62L^+^CD44^−^, CD62L^−^CD44^+^, and CD62L^+^CD44^+^ subsets within conventional (Foxp3^−^) CD4^+^ T cells from the LNs of naïve mice. **(B)** Superimposed bar charts illustrate the frequencies of YFP^+^ cells in total conventional CD4^+^ and CD8^+^ T cells of naïve mice and mice with EAE. The superimposed bar charts illustrate frequencies of YFP^+^ cells in CD62L^−^CD44^−^, CD62L^+^CD44^−^, CD62L^−^CD44^+^, and CD62L^+^CD44^+^ subsets of conventional CD4^+^ (upper) and CD8^+^ (lower) T cells from **(C)** naïve mice and **(D)** mice with EAE. Data represents two independent experiments with naïve (total n = 7) and mice with EAE (total n = 8). Error bars indicate the mean ± SEM. Statistical analysis was conducted using the Student’s t-test; *P<0.05.

An unexpected finding was that 10-20% of naïve CD4□ cells (CD62L CD44) were YFP^+^, indicating that these cells expressed GM-CSF. This was surprising given that naïve T cells do not express cytokines, including GM-CSF. We hypothesized that these YFP^+^ naïve CD4^+^ T cells could be stem cell memory T cells (TSCM), so we stained splenocytes of naïve mice for TSCM markers, including Sca-1, and CD122, as well as other T cell subset markers such as CD95, CD27, and CD127. However, we did not find differences between YFP□ and YFP□ naïve CD4^+^ T cell populations (**Figs. S4A and B**). Additionally, no significant differences were found in current GM-CSF, IFN-γ, IL-17, or IL-2 expression between these populations following stimulation with PMA and Ionomycin (**Fig. S4C**), suggesting that YFP^+^ CD4^+^CD62^+^CD44^−^ T cells are genuinely naïve. It is unclear what led these cells to express GM-CSF and if some naïve T cells also express other cytokines.

### T cells simultaneously express both GM-CSF and CXCR6

CXCR6 plays a critical role in T cell trafficking and retention within tissues, particularly in the lungs, liver (20, 21), and CNS of mice with EAE (22). Recent studies have suggested a link between CXCR6 expression and GM-CSF production in T cells, especially CD4^+^ T cells, under specific inflammatory conditions or within certain tissue microenvironments (10). To investigate the correlation between GM-CSF and CXCR6 expression, we analyzed YFP and CXCR6 expression across various T cell subsets in steady state and CNS inflammation. Conventional (Foxp3^−^) CD4^+^YFP^+^CXCR6^+^ T cell frequencies were higher in the liver compared to other organs of naïve mice, with no significant changes observed in mice with EAE. However, this population increased in the spleen, lung, and CNS of mice with EAE. CD8^+^YFP^+^CXCR6^+^ cells had the highest frequency in the SI, followed by the CNS and liver, with an increase in the lungs of mice with EAE (**Figs. 4A and B**). The frequencies of CXCR6^+^ cells among CD4^+^YFP^+^ and CD8^+^YFP^+^ cells were several times higher than among their YFP^−^ counterparts, except in the case of CD8^+^ T cells from the SI. Approximately 20% of CD8^+^YFP^+^ and CD8^+^YFP^−^ cells expressed CXCR6, regardless of immunization status (**Figs. 4C and D**). Notably, CD4^+^ and CD8^+^ T cells from the liver and CNS exhibited the highest levels of CXCR6 expression among YFP^+^ cells (**Figs. 4A, C, and D**).

**Figure 4.**
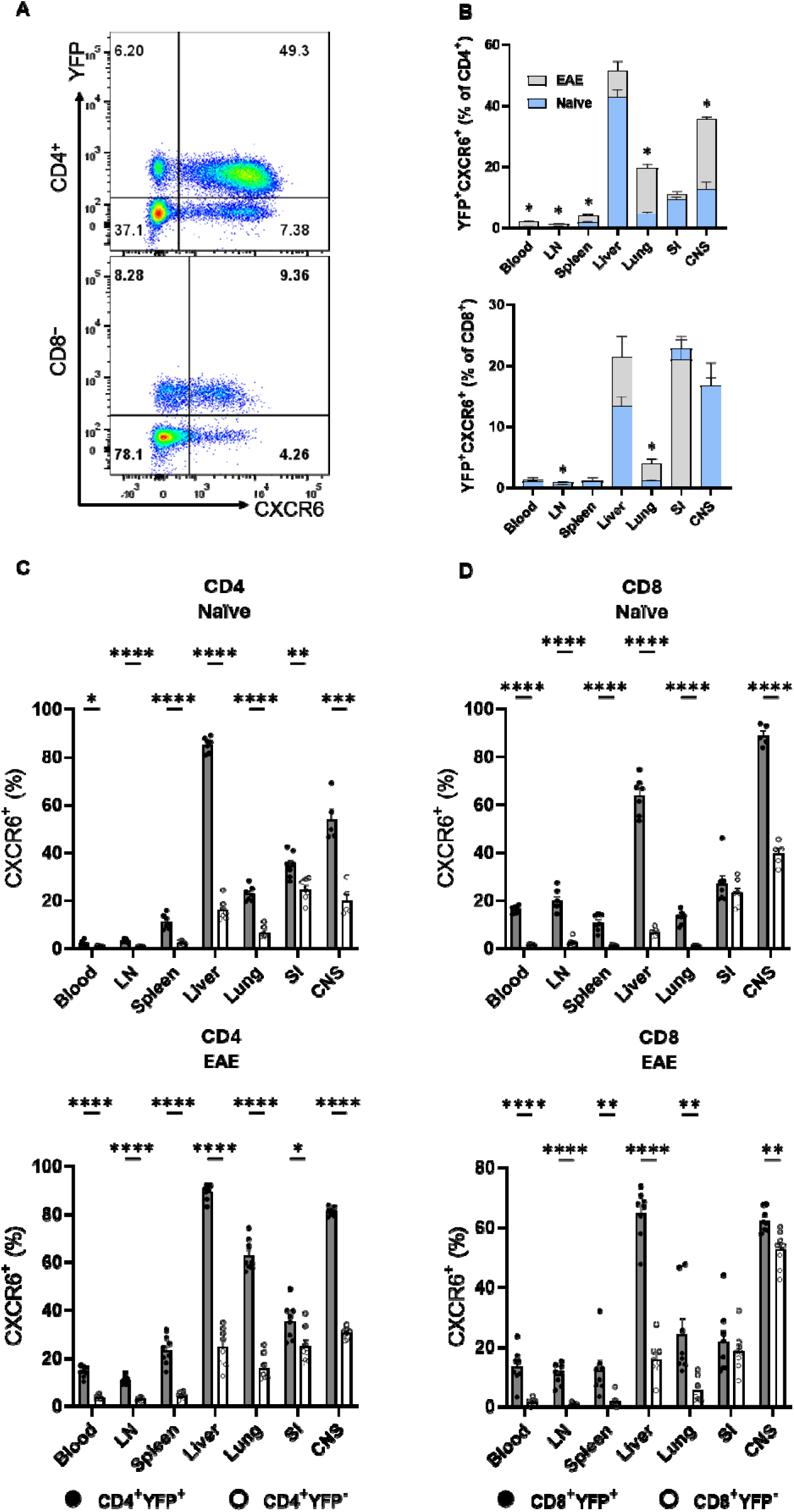
YFP and CXCR6 expression by CD4^+^ and CD8^+^ T cells. Mononuclear cells were isolated from the blood, LNs, spleen, liver, lung, SI, and CNS of 2-3-month-old male and female Gr/fr mice. The cells were stained for CD45, CD4, CD8, CD62L, CD44, and CXCR6, and evaluated for YFP expression within different subsets. **(A)**, Representative flow cytometry dot plots show YFP and CXCR6 expression by gated conventional (FoxP3^−^) CD4^+^ (upper panel) and CD8^+^ (lower panel) T cells from the liver of naïve mice. **(B)**, Superimposed bar charts for YFP^+^CXCR6^+^ cells of gated CD4^+^ (upper panel) and CD8^+^ (lower panel) T cells of naïve and mice with EAE. **(C)**, Percentages of YFP^+^CXCR6^+^ and **(D)**, YFP^−^CXCR6^−^ CD4^+^, and CD8^+^ T cells from naïve (upper panels) and mice with EAE (lower panels). Data represents two independent experiments with naïve (total n = 7) and mice with EAE (total n = 8). Error bars indicate the mean ± SEM. Statistical analysis was conducted using the Student’s t-test; *P<0.05; **P<0.01; ***P<0.001; ****P<0.0001. Abbreviations: LN- lymph nodes, SI-small intestine, CNS- central nervous system.

While 10-20% of naïve (CD62L^+^CD44^−^) CD4^+^ T cells expressed YFP, this subset was vastly CXCR6^−^. In contrast, CD4^+^CD62L^+^CD44^+^YFP^+^ and CD4^+^CD62L^−^CD44^+^YFP^+^ subsets had the highest proportions of CXCR6^+^ cells across all organs in both naïve mice (**Fig. S5A**) and those with EAE (**Fig. S5B**). Our findings highlight a significant association between CXCR6 and YFP expression in T cells, except for naïve T cells. During EAE, there was almost 100% overlap between YFP and CXCR6 expression in some organs, such as the liver, lung, and CNS. We also observed similarly pronounced co-expression of YFP, CXCR3, and CXCR6 (**Figs. S5C and D**). Conflicting reports in the literature suggest that CXCR6 and CXCR3 either contribute to EAE development or have no impact (22, 23). In our study, neither CXCR3 nor CXCR6 played an important role in EAE (**Fig. S5E**).

### A subset of CD4^+^Foxp3^+^ T cells express GM-CSF

CD4^+^Foxp3^+^ regulatory T cells (Tregs) play crucial roles in maintaining immune homeostasis across various organs (24). While Tregs primarily function to suppress immune responses, they can also produce cytokines associated with Th lineages, such as IFN-γ and IL-17, particularly under inflammatory conditions (25, 26). We noticed that some CD4^+^Foxp3^+^ cells were YFP^+^, indicating that they produce GM-CSF. Given that GM-CSF production by Treg cells has not been previously described, we further characterized their GM-CSF expression.

In naïve mice, 8-18% of CD4^+^ T cells across various organs were Foxp3^+^. This percentage increased following immunization, particularly in the CNS (**Fig. 5A**). In both naïve and EAE mice, approximately 10% of CD4^+^Foxp3^+^ T cells were YFP^+^ (**Fig. 5B**), indicating a history of GM-CSF expression. We analyzed cytokine and YFP expression in CD4^+^Foxp3^+^ cells from the spleen of naïve and EAE mice. A small percentage of YFP^+^ cells from naïve mice and a larger percentage in EAE mice expressed GM-CSF, IFN-γ, and IL-17 at significantly higher levels compared to YFP^−^ cells (**Fig. 5C**). Additionally, we evaluated the expression of Helios and neuropilin-1 (CD304), two markers predominantly associated with thymic Treg (tTreg) and peripheral Treg (pTreg) (27). Similar proportions of YFP^+^ and YFP^−^ cells expressed both markers (**Fig. 5D**). In conclusion, CD4^+^Foxp3^+^ T cells expressing YFP display increased inflammatory cytokine production, suggesting a higher phenotype similarity with conventional CD4^+^ T cells than its YFP^−^ counterparts.

**Figure 5.**
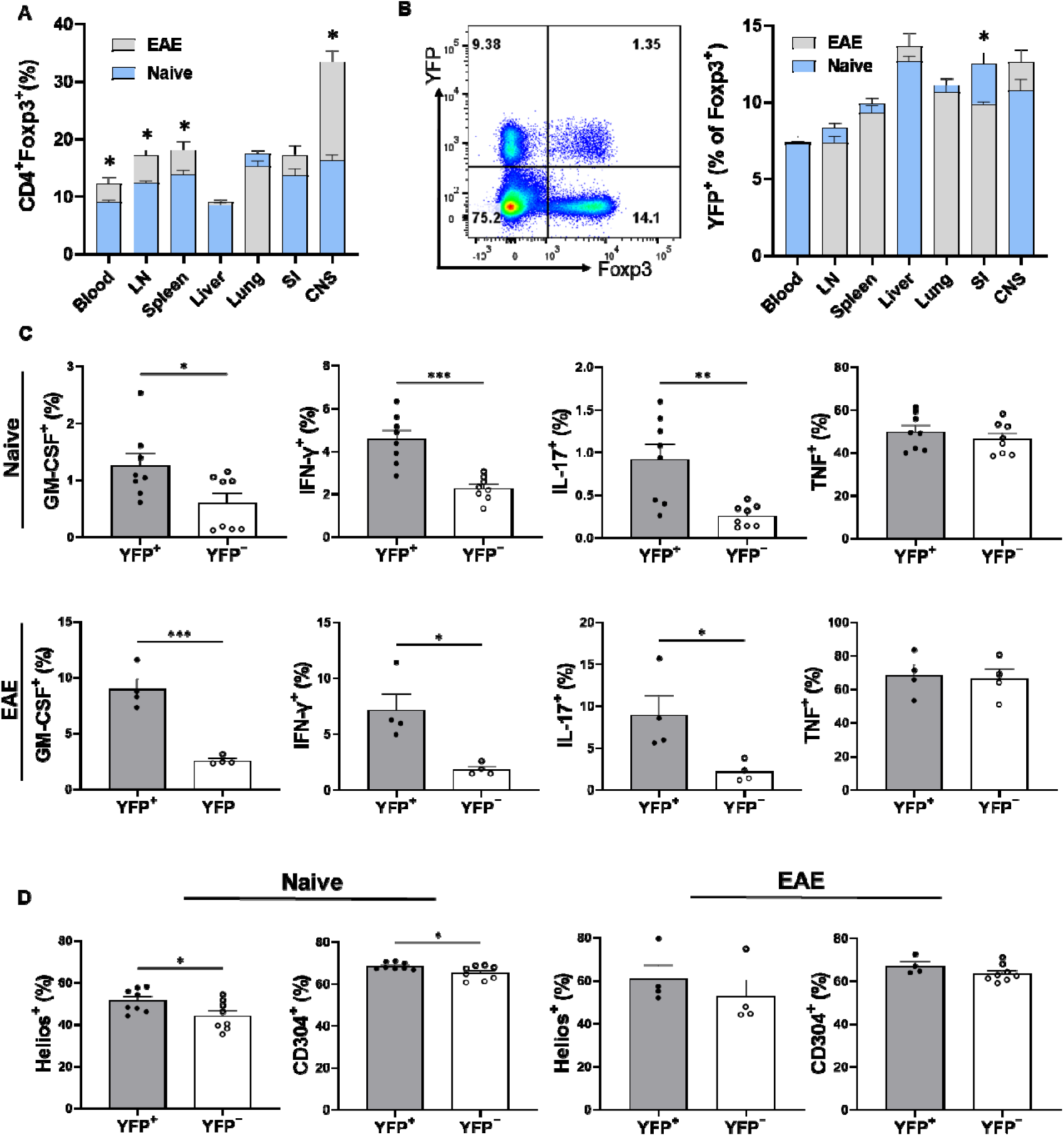
YFP and cytokine expression by Foxp3□CD4□ T cells. Mononuclear cells were isolated from the blood, LNs, spleen, liver, lung, SI, and CNS of 2- to 3-month-old male and female Gr/fr mice, both naïve and with EAE. The cells were stained for CD45, CD4, and Foxp3, and assessed for YFP expression by flow cytometry. **(A)** The superimposed bar chart shows the frequency of Foxp3□ cells among CD4□ T cells in various organs of naïve mice and mice with EAE. **(B)** Representative flow cytometry dot plots show YFP and Foxp3 expression by gate CD4^+^ T cells from the spleens of naïve mice. The superimposed bar chart shows the frequencies of YFP^+^CD4□Foxp3□ cells from various organs of naïve and EAE mice. **(C)** Total CD4□ T cells were isolated from the spleens of naïve and EAE mice (20 dpi.) using magnetic-activated cell sorting (MACS) and stimulated with PMA/Ionomycin for 5 h. Foxp3□YFP□CD4□ and Foxp3□YFP□CD4□ cells were evaluated for GM-CSF, IFN-γ, IL-17, and TNF expression. **(D)** Expression of Helios and CD304 by Foxp3□CD4□ T cells. Data represents two independent experiments (total n = 7–8 for naïve mice and n = 8 for mice with EAE, except for panels C and D, where n = 4 for EAE). Error bars indicate mean ± SEM. Statistical analysis was performed using Student’s t-test; *P < 0.05, **P < 0.01, ***P < 0.001. Abbreviations: LN- lymph nodes, SI - small intestine, CNS- central nervous system.

### A significant correlation exists between current cytokine production and the history of GM-CSF expression in T cells

Across various organs of both naïve and EAE mice, 10-30% of conventional CD4^+^ T cells and 10-70% of CD8^+^ T cells were YFP^+^. To assess cytokine production by YFP^+^ cells, we stimulated MNCs with PMA/Ionomycin and analyzed the expression of several cytokines by flow cytometry. Overall, except for TNF, the expressions of GM-CSF, IFN-γ, IL-17, IL-4, and IL-10 were higher in non-lymphoid organs (liver, lung, and SI) and the CNS compared to blood and lymphoid organs of both naïve and EAE mice. In general, a greater proportion of CD4^+^ T cells from mice with EAE-expressed cytokines. Following EAE induction, CD8^+^ T cells showed an upregulation in IL-17, IL-4, and IL-10 expression (**Fig. S6**). Comparison of cytokine expression between CD4^+^ YFP^+^ and YFP^−^ cells revealed a marked tendency of YFP^+^ cells from naïve and EAE mice to produce cytokines, particularly GM-CSF, IL-17, and IFN-γ (**Fig. S7**). This was not surprising, as the frequency of cells with an effector memory phenotype (CD44) was higher among YFP cells (∼27% in the spleen of naïve mice) compared to YFP cells (∼17% in the spleen of naïve mice), with the difference being approximately two-fold higher in non-lymphoid organs. In the CD8^+^ cells of both naïve and EAE mice, a markedly bigger proportion of YFP^+^ cells produced GM-CSF and IFN-γ compared to YFP^−^ cells (**Fig. S8**). We found a significant association between YFP expression and the production of GM-CSF, IFN-γ, and IL-17 by CD4^+^ T cells from the liver, lung, and CNS of naïve mice, with a notable increase in their frequency in EAE mice (**Figs. 6A and B**). Overall, CD4^+^YFP^+^ T cells constitute a multi-cytokine-producing subset primarily characterized by the expression of GM-CSF, IFN-γ, TNF, and IL-17, with minimal expression of IL-4 and IL-10. A similar increase of IL-17 expression by CD8^+^YFP^+^ cells was noted after immunization for EAE induction (**Fig. 6C**).

**Figure 6.**
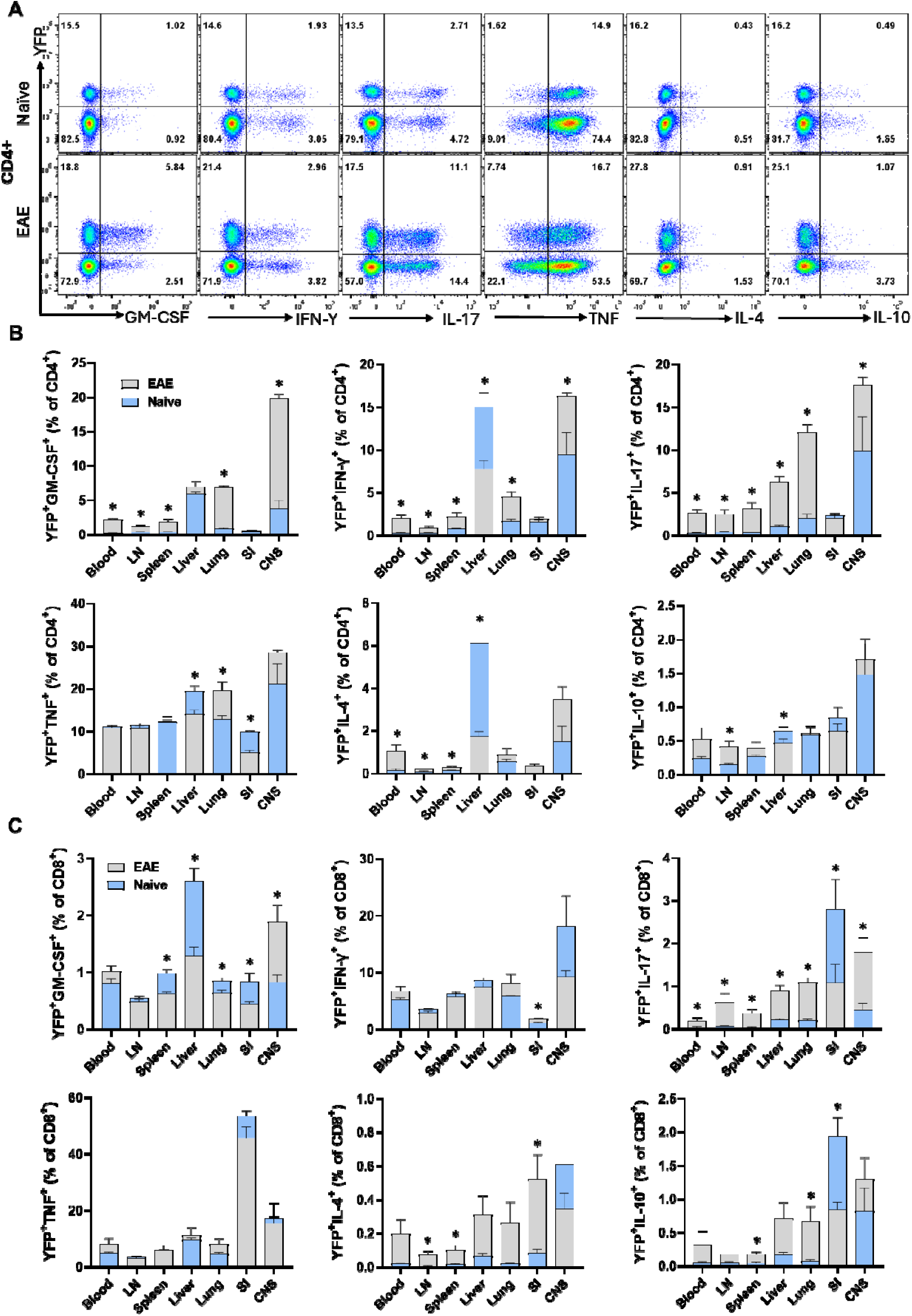
YFP and cytokine expression by CD4^+^ and CD8^+^ T cells. Mononuclear cells were isolated from the blood, LNs, spleen, liver, lung, SI, and CNS of 2–3-month-old male and female naïve and EAE Gr/fr mice. Cells were stimulated with PMA, Ionomycin, and GolgiPlug and stained for surface and intracellular antigens. **(A)**, Flow cytometry plots illustrate gating for YFP and GM-CSF, IFN-γ, IL-17, TNF, IL-4, and IL-10 in gated lung CD4^+^ T cells of naïve and mice with EAE. Superimposed bar charts illustrate YFP and cytokines GM-CSF, IFN-γ, IL-17, TNF, IL-4, and IL-10 expression by gated CD4^+^ **(B)** or CD8^+^ **(C)** T cells. Results are expressed as the mean ± SEM; naïve (total n = 7) and mice with EAE (total n = 8) from 2 independent experiments. Statistical analysis was performed using Student’s t-test; *P<0.05. Abbreviations: LN- lymph nodes, SI- small intestine, CNS- central nervous system.

CXCR5, a marker for T follicular helper (TFH) cells, is involved in T cell recruitment in lymphoid tissues (28). The YFP^+^CXCR5^+^ population constituted a small proportion of CD4^+^ and CD8^+^ T cells (less than 5%) across different organs of naïve mice. Following immunization, there was a variable change in this population among CD4^+^ T cells and a decrease among CD8^+^ T cells (**Fig. S9**).

### GM-CSF expression is a transient characteristic of CD4□ T cells under steady-state conditions

Our data show that only a portion (2-18%) of CD4□YFP□T cells from the organs of naïve mice produced GM-CSF upon ex vivo stimulation with PMA and Ionomycin, while the portion of YFP cells that expressed GM-CSF was even smaller (1-5%) (**Fig. S7A**). These data indicate that GM-CSF expression is transient and likely permanently ceases in almost all the cells that used to express it. Furthermore, ThGM and Th1 cells that developed in vitro started to lose capacity for GM-CSF production after 72 h (**Fig. S10)**. However, we hypothesized that short-term ex vivo stimulation with PMA and Ionomycin may not reflect the behavior of cells upon activation by their T cell receptor (TCR) over several days. To investigate this in vivo within the context of EAE, we analyzed GM-CSF and YFP expressions in the CNS at disease onset (15 d.p.i.), peak (20 d.p.i.), and beginning of chronic phase (27 d.p.i.). The frequency of GM-CSF^+^CD4^+^ T cells decreased from 32% at the onset to 18% at the chronic phase. Conversely, YFP^+^GM-CSF^−^ cells increased from 21% at the onset to 28% during the chronic phase (**Fig. 7**).

**Figure 7.**
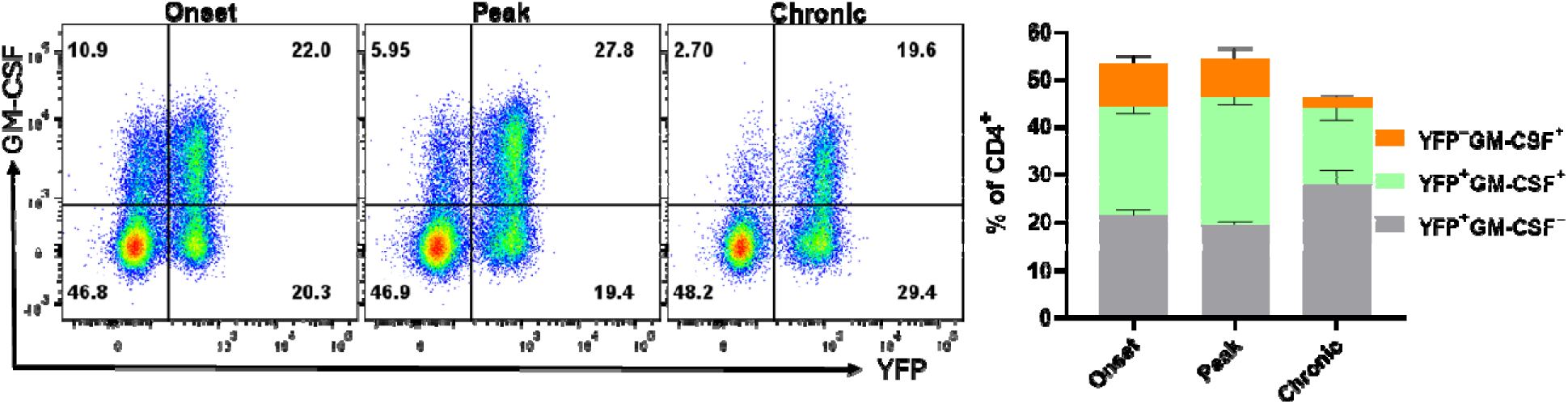
YFP and GM-CSF expression in CD4^+^ T cells from the CNS of EAE mice. Mononuclear cells were isolated from the CNSs of 2–3-month-old male and female Gr/fr mice during the onset (15 dpi), peak (20 dpi), and chronic (27 dpi) phases of EAE. Cells were stimulated with PMA, Ionomycin, and GolgiPlug for 6 h, followed b staining for surface and intracellular antigens. Flow cytometry plots illustrate the gated YFP and GM-CSF expression within gated CD4^+^ T cells among live CD45^hi^ cells. Results for the YFP^−^GM-CSF^+^, YFP^+^GM-CSF^+^, and YFP^+^GM-CSF^−^ populations are presented as mean ± SEM by the stacked bar chart. Data represents mice with EAE (n = 4 per group) from one experiment.

To test if CD4□YFP□T cells restart their GM-CSF expression upon activation through their TCR, we sorted YFP^+^ and YFP^−^ CD4□CD25□ T cells from the spleen, liver, and lungs. These cells were activated with anti-CD3/CD28 antibodies in co-culture with T cell-depleted splenocytes for 3 days. Following stimulation, flow cytometry analysis revealed a large increase in GM-CSF^+^ cells, reaching 10-20% of both YFP^+^ and YFP^−^ CD4^+^ T cells. The proportions of IFN-γ^+^ cells also markedly increased, reaching 20-60% in both YFP^+^ and YFP^−^ subpopulations, whereas the proportions of IL-17^+^ cells notably declined, presumably because of the switch from Th17 to Th1 (**Figs. 8A-C**). Measuring cytokine concentrations in the cell culture supernatants showed that both YFP^+^ and YFP^−^ cells from all three organs secreted similar amounts of GM-CSF and IFN-γ, and only the concentration of IL-17 was significantly higher in the YFP culture compared to the YFP□ culture (**Fig. 8D**). Although 10-20% of CD4□YFP□ cells expressed GM-CSF, only about 7% of these cells became YFP 72 h after activation (**Fig. 8E**), suggesting that switching on YFP expression may lag behind GM-CSF production. However, it is possible that not all cells that were GM-CSF^+^ upon PMA and Ionomycin stimulation expressed GM-CSF during the preceding 3-day activation period. The capacity of cells to produce cytokine upon PMA and Ionomycin stimulation does not always correlate with the extent of prior cytokine production. Hence, cells that could produce GM-CSF but didn’t or only did so in small quantities or briefly are unlikely to become YFP^+^ because only a small amount of Cre, if any, was produced, rendering the activation of YFP expression unlikely.

**Figure 8.**
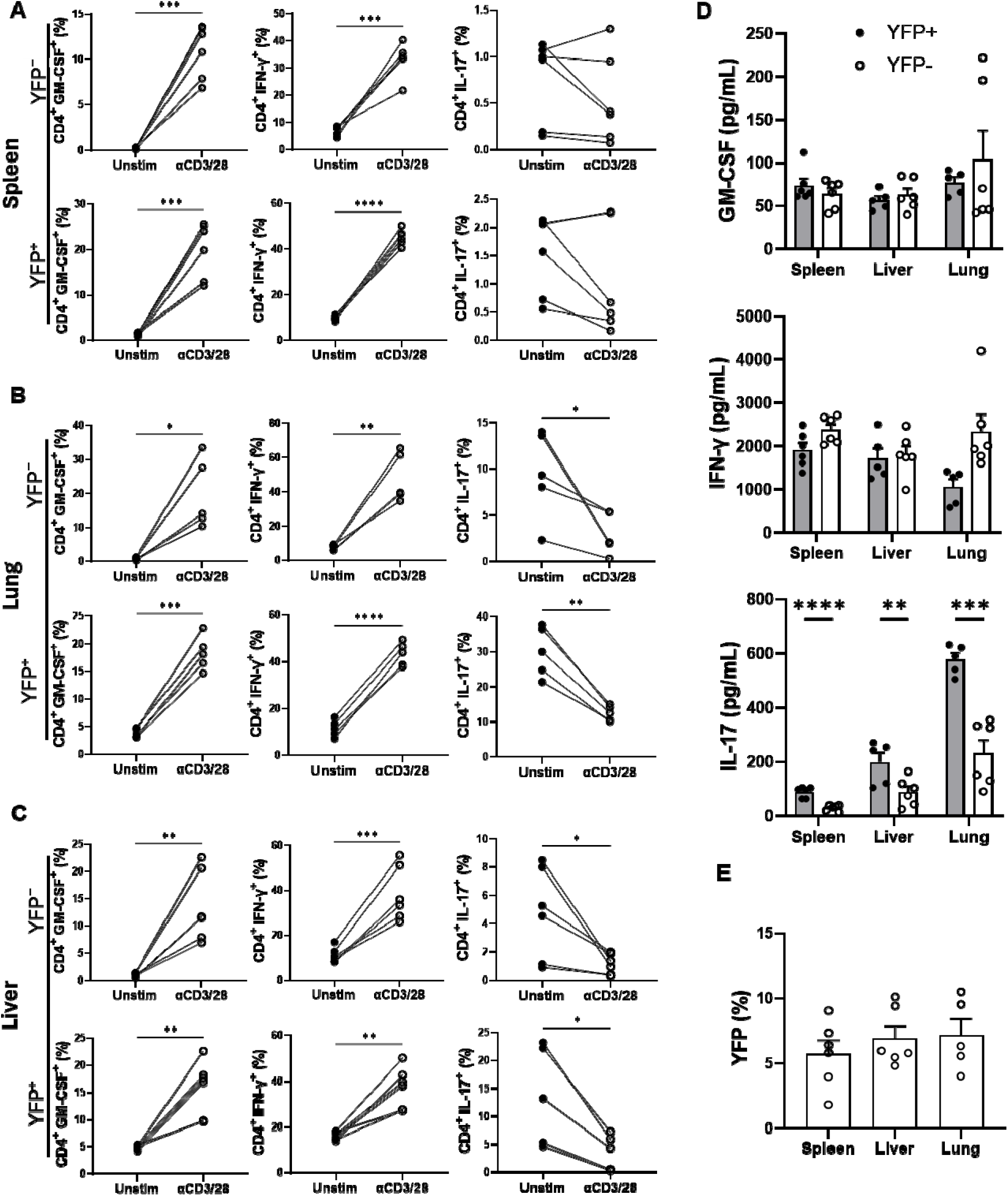
GM-CSF, IFN-γ, and IL-17 expression by YFP□ and YFP^−^ CD4□ T cells following TCR activation. YFP□ and YFP^-^ CD25□ CD4□ cells were sorted from the spleen, liver, and lung mononuclear cells of naïve 2-3-month-old male and female Gr/fr mice. The sorted cells were either directly stimulated with PMA, Ionomycin, an GolgiPlug, or co-cultured for 72 h with T cell-depleted splenocytes, and with or without soluble anti-CD3/CD2 mAbs (3 µg/ml); cells were then stimulated with PMA, Ionomycin, and GolgiPlug and stained for CD4, GM-CSF, IFN-γ, and IL-17. Cytokine expression was analyzed in gated YFP□CD4□ and YFP□CD4□ cells from the spleen **(A)**, lung **(B)**, and liver **(C)**, with or without anti-CD3/CD28 mAb stimulation. Results were analyzed using paired Student’s t-test. **(D)** Cytokine production by YFP□CD4□ and YFP□CD4□ cell culture following anti-CD3/CD28 mAbs stimulation. **(E)** The proportion of YFP^−^CD4□ cells switching to YFP□CD4□ following 72 h activation in culture. Data is shown as the mean ± SEM from 3 independent experiments. Statistical significance was determined using the Paired t-test (A, B, C) and unpaired Student’s t-test (for D and E). *P<0.05; **P<0.01; ***P<0.001; ****P<0.0001.

In the next step, we conducted a similar experiment, but with sorted YFP^+^ and YFP^−^ naïve and effector/memory CD4^+^ T cells from the spleen of naïve mice. Following activation with anti-CD3/28 mAbs, both YFP^+^ and YFP^−^ naïve CD4^+^ T cells similarly expressed GM-CSF (25-30%), whereas both YFP^+^ and YFP^−^ effector memory CD4^+^ T cells barely expressed GM-CSF (2-5%) (**Figs. 9A and B**). GM-CSF concentrations in culture supernatants agreed with flow cytometry data (**Fig. 9C**). The percentages of IFN-γ^+^ and IL-17^+^ cells were similar among all four tested populations. Interestingly, both YFP^+^ and YFP^−^ subsets of effector memory CD4^+^ T cells secreted notable amounts of IL-17 in culture supernatants, while naïve cells secreted almost none (**Fig. 9C**). In addition, the rate of naïve CD4^+^YFP^−^ T cells converting to YFP^+^ was greater than that of effector memory CD4^+^YFP^−^ T cells (**Fig. 9D**). It is surprising that effector memory cells, even those that previously expressed GM-CSF, did not re-express it upon activation. This suggests that the increase in GM-CSF^+^ cells after activation among total YFP^+^ and YFP^−^ cells shown in **Fig. 8A** is due to de novo upregulation of GM-CSF expression in naïve cells. However, we noticed that effector memory cells in our cultures rapidly declined after activation and that only a fraction survived for three days. Hence, it is possible that effector memory cells that produced GM-CSF already died by the time of analyses and are therefore not included in our data. However, when splenocytes of naïve mice are directly ex vivo stimulated with PMA and Ionomycin, there were almost no GM-CSF^+^ cells among YFP^+^CD4□ T cells (**Fig. 6B**), suggesting that effector memory YFP^+^CD4^+^ T cells indeed do not express GM-CSF.

**Figure 9.**
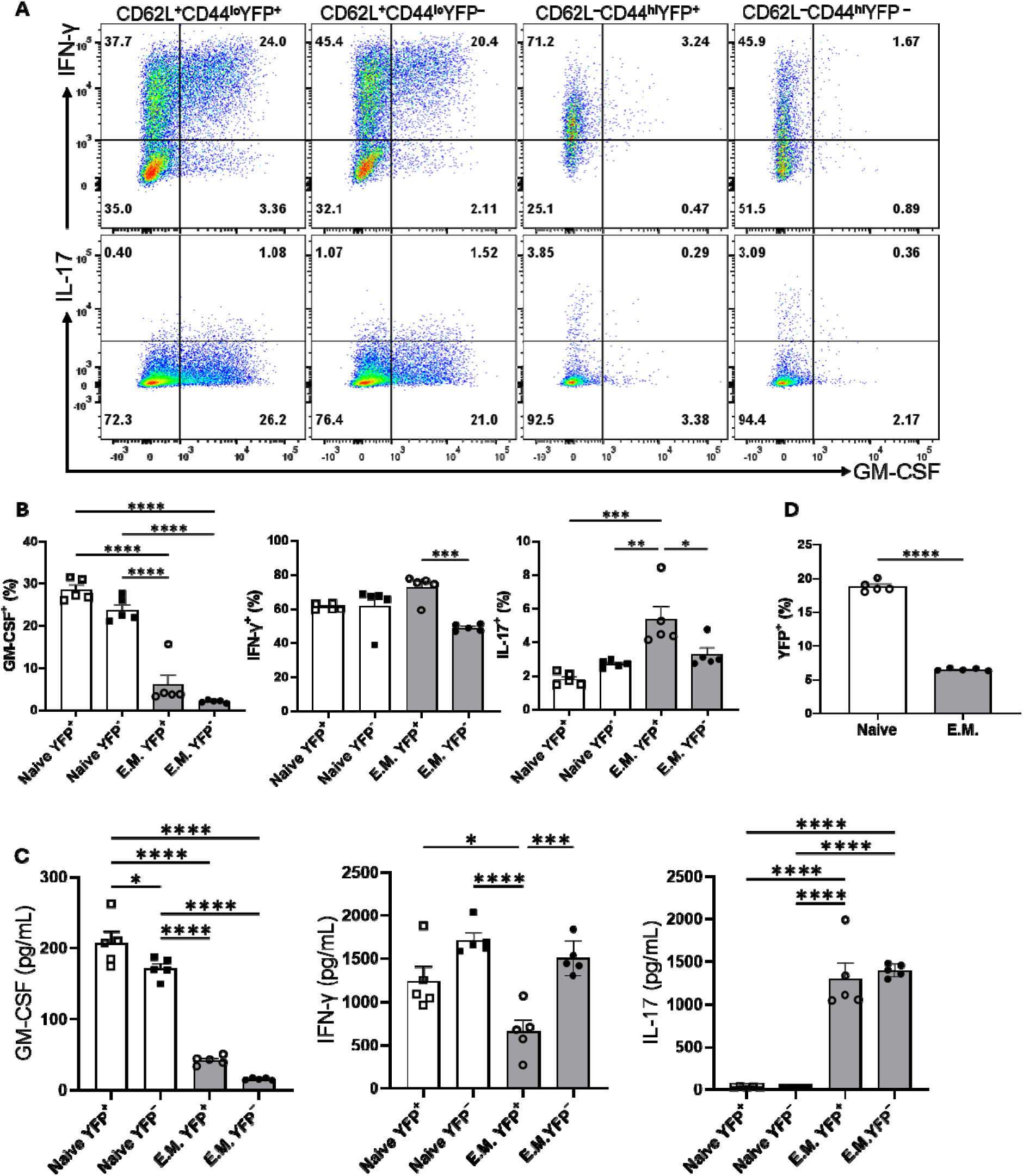
GM-CSF, IFN-γ, and IL-17 expression by YFP□ and YFP^−^ populations of naïve and effector memory CD4□ T cells following stimulation. Conventional naïve (CD25^−^CD62L□CD44^lo^CD4□) YFP□ an YFP^−^, and effector memory (CD25^−^CD62L□CD44^hi^CD4□) YFP□ and YFP^−^ T cells were sorted from the spleens of naïve 2-3-month-old male and female Gr/fr mice. The cells were co-cultured with T cell-depleted splenocytes and soluble anti-CD3/CD28 mAbs (3 µg/ml) for 72 h, then stimulated with PMA, Ionomycin, and GolgiPlug. Cells wer stained against CD4, GM-CSF, IFN-γ, and IL-17. **(A)** Flow cytometry plots show gating for GM-CSF, IFN-γ, and IL-17 in gated CD4□ T cells. **(B)** Cytokine expression in gated CD4□ T cells from cultured cells following anti-CD3/CD28 mAbs stimulation. **(C)** Cytokine levels were measured by ELISA in the culture supernatants after anti-CD3/CD28 mAbs stimulation. **(D)** The proportions of YFP^+^ cells in naïve and effector memory YFP^−^ populations after 72 h of activation and co-culture. Data are presented as the mean ± SEM from 2 independent experiments (total n=5). Statistical analysis was conducted using an unpaired Student’s t-test and one-way ANOVA. *P < 0.05; **P < 0.01; ***P < 0.001; ****P < 0.0001.

### The transcriptomes of YFP^+^ and YFP**^−^** naïve CD4^+^ T cells are highly similar, whereas those of YFP^+^ and YFP**^−^**CD4^+^ effector memory T cells are markedly distinct

To identify differences between CD4^+^ T cells that did and did not produce GM-CSF^+^, we sorted YFP^+^ and YFP^−^ naïve and CD4^+^ effector memory T cells from the spleens of naïve *Gr/fr* mice and analyzed their transcriptomes using bulk RNA-seq. Comparison between CD4^+^ effector memory subpopulations identified 262 genes differentially upregulated in YFP^+^ and 652 genes in YFP^−^ cells (**Fig. 10A**), while only a few genes were differentially expressed between YFP^+^ and YFP^−^ naïve CD4^+^ T cells (**Fig. 10B**). YFP^+^ CD4^+^ effector memory T cells showed higher expression of *Csf2*, *Il4*, *Rorc*, *Tbx21*, and *Cxcr6* with lower expression of *Foxp3* (**Figs. 10A and C**). On the other hand, YFP^+^ naïve CD4^+^ T cells only showed higher expression of *Chn2* and *Crybg3* and lower expression of *Cd8b1* compared to YFP^+^ naïve CD4^+^ T cells (**Fig. 10B**). We next tested some of these findings from the steady state in mice with EAE. In agreement with RNA-seq results, most YFP^+^ CD4^+^ effector memory T cells from the CNS of mice with EAE expressed GM-CSF. They also displayed higher expression of *T-bet* and lower expression of *Foxp3* (**Fig. 10D**). Unlike YFP^+^ CD4^+^ effector memory T cells in a steady state, these EAE cells had higher frequencies of IL-17^+^, IFN-γ^+^, TNF^+^, and IL-2^+^ cells and higher expression of *GATA3* (**Fig. 10D**). Collectively, these results show that YFP^+^ and YFP^-^ naïve CD4^+^ T cells are virtually indistinguishable, whereas YFP^+^ and YFP^-^ CD4^+^ effector memory T cells differ to a surprisingly large extent, both in a steady state and inflammation.

**Figure 10.**
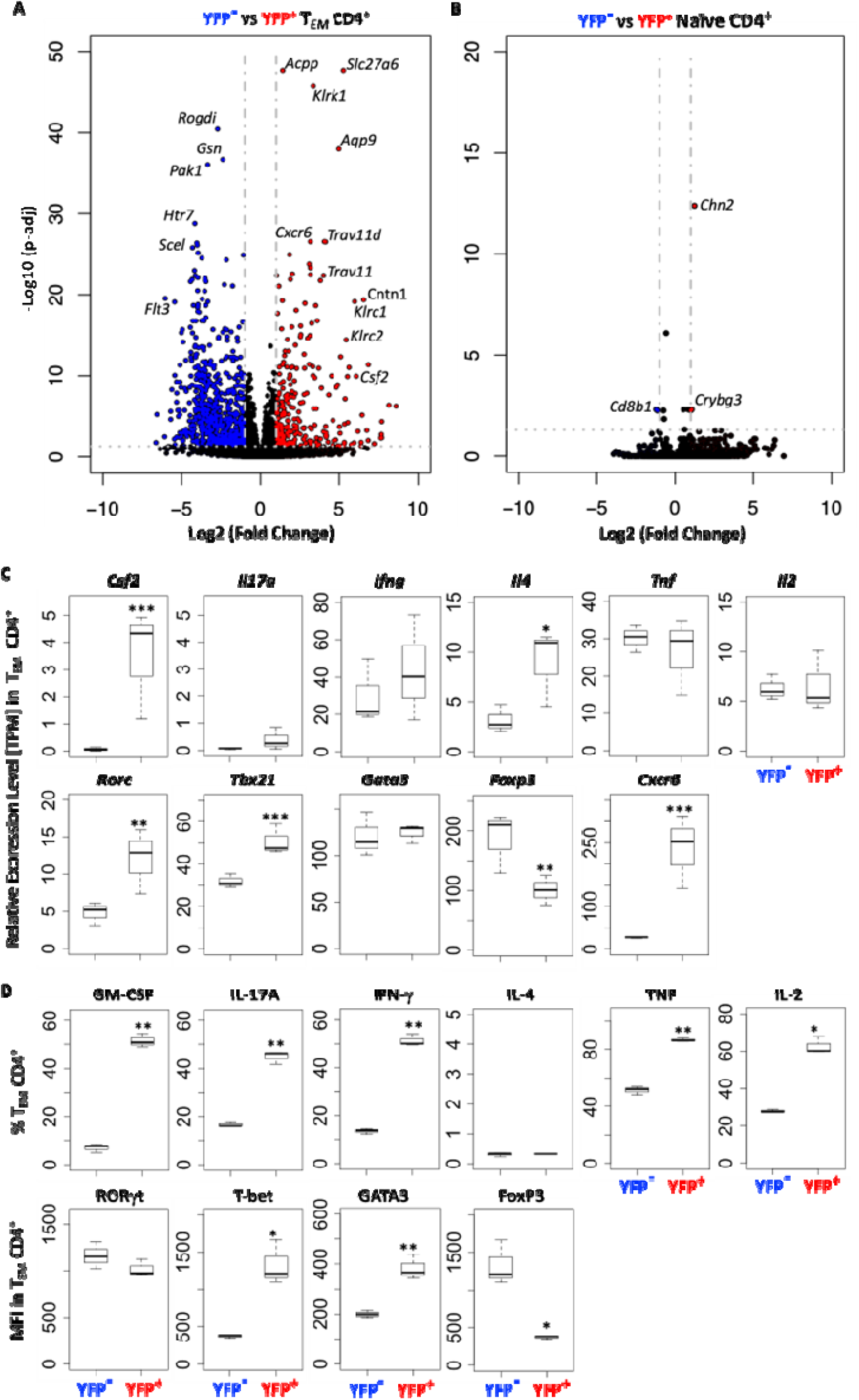
YFP^+^ and YFP^-^ CD4^+^ T_EM_ cells have a distinct transcriptomic profile. YFP^−^ and YFP^+^ naïve (CD62L0CD44^lo^) and T_EM_ (CD62L^−^CD44^hi^) CD4□ T cells were FACS sorted from spleens of naïve *Gr/fr* mice (n=3), RNA was isolated and analyzed by RNA-seq. **(A)** Volcano plot shows 914 differentially expressed genes between YFP^+^ and YFP^−^ CD4□ T_EM_ cells, with 262 genes upregulated in YFP^+^ and 652 genes in YFP^−^ cells. **(B)** Volcano plot shows 3 differentially expressed genes between YFP^+^ and YFP^−^ naïve CD4□ T cells, with 2 genes upregulated in YFP^+^ and 1 gene in YFP^−^ cells. **(C)** Normalized gene expression levels (TPM) of several markers expressed by YFP^+^ and YFP^−^ CD4^+^ T_EM_ cells from RNA-seq analysis. **(D)** YFP^+^ and YFP^−^ CD4^+^ T_EM_ cells from the CNS of *Gr/fr* mice (n=3) immunized for EAE induction were analyzed at 18 d.p.i. Box plots show frequencies of GM-CSF^+^, IL-17A^+^, IFN-g^+^, IL-4^+^, TNF^+^, and IL-2^+^ YFP^+^ and YFP^−^ CD4^+^ T_EM_ cells in the CNS of mice with EAE. The bottom row box plot shows the staining intensity (MFI) of RORγt, T-bet, GATA3, and Foxp3 in YFP^+^ and YFP^−^ CD4^+^ T_EM_ cells in the CNS of mice with EAE. Box plots show interquartile range (IQR), with horizontal lines denoting median. P-values were calculated using paired Student’s t-test; * p < 0.05, ** p < 0.01, *** p < 0.001.

## Discussion

GM-CSF is primarily viewed as a key pro-inflammatory cytokine that elicits the inflammatory phenotype in myeloid cells. GM-CSF is produced at relatively low levels in a steady state, but its production significantly increases during immune responses. Various immune cell types produce GM-CSF, with T cells being its primary source during inflammation (10, 11, 17, 29). Given its essential role in myeloid cell function during inflammation, including autoimmunity, the biology of GM-CSF has been studied extensively and from various angles. However, the relationship between past and ongoing GM-CSF expression by diverse immune cells across different organs is only partially understood. In this study, we took advantage of our GM-CSF fate mapping mouse line to trace the lineage and function of GM-CSF-expressing cells in steady state and autoimmune neuroinflammation. Overall, we found that the capacity to express GM-CSF is transient, as the vast majority of cells that previously expressed it at some point do not express it again, at least not upon stimulation ex vivo. Unexpectedly, myeloid cells, primarily macrophages, were the dominant GM-CSF-producing cell type in some organs, such as the liver, lung, and spleen. In other organs, such as lymph nodes, small intestine, and inflamed CNS, T cells were the leading producers of GM-CSF. Among CD4^+^ T cells, most GM-CSF-producing cells were found in the effector memory subset, as expected. Surprisingly, subsets of naïve CD4□ T and Tregs also exhibited a history of GM-CSF expression. Prior GM-CSF expression by effector memory CD4^+^ T cells strongly correlated with a distinct transcriptomic profile. We also found that GM-CSF-producing T cells co-express CXCR6, suggesting a link to tissue residency.

In both naïve and EAE mice, CD4□ T cells, γδ T cells, NK cells, and CD11b cells display the highest GM-CSF expression (primarily past GM-CSF expression). In contrast, B cells were unique in predominantly expressing GM-CSF actively, albeit at low levels. This may suggest that GM-CSF production in B cells may be an activation or differentiation feature rather than a general characteristic under normal conditions (16). Consistent with our findings, Komuczki et al. reported that active GM-CSF expression is nearly undetectable before EAE induction but emerges in LNs at disease onset, primarily in CD4□ and CD8□ T cells, γδ T cells, and NK cells. Notably, most of these cells had a history of GM-CSF expression, while only a subset actively produced it (10). Given that different immune cells express GM-CSF at varying levels and their frequencies differ across organs, we focused on the YFP population to determine the primary GM-CSF sources in each organ. Among CD45□ YFP□ cells, the dominant populations varied by tissue. In the blood, spleen, and lungs of naïve mice, CD11b cells were the predominant YFP population, followed by CD4□ T cells. In LNs and the thymus, CD4□ T cells were the major YFP cells, whereas, in the liver, both CD11b cells and CD4□ T cells were prominent. In the SI, γδ T cells and CD8□ T cells were the primary YFP populations. Although the distribution of GM-CSF-expressing cells largely reflects the overall frequency of immune cell subsets across tissues, our findings suggest that CD4□ T cells and CD11b cells (primarily macrophages) are the predominant sources of GM-CSF in mice bodies. While these cells are the main GM-CSF producers under steady-state conditions, the disease context and tissue microenvironment critically regulate its expression. Consistent with findings from Komuczki et al. (10) and Sheng et al. (30), our results show that in the inflamed CNS, effector CD4□ T cells are the dominant contributors to both past and current GM-CSF production, with smaller contributions from CD8□ T cells, NK cells, γδ T cells, and CD11b cells. In contrast, in inflamed joints during inflammatory arthritis, NK cells and macrophages are the primary GM-CSF producers (29, 31), while patients with spondyloarthritis exhibit increased GM-CSF-producing CD4 and CD8 lymphocytes in their joints, along with an expansion of GM-CSF-producing type 3 IL3 (ILC3s) in synovial tissues (32). ILC3s also play a key role in GM-CSF production during intestinal inflammation (17). These findings highlight the complex interplay between immune cell type, organ-specific microenvironments, and inflammatory conditions in shaping GM-CSF expression.

In our study, 10-20% of naïve CD4□ T cells (but not naïve CD8□ T cells) were YFP, indicating prior GM-CSF expression. This was unexpected, as naïve T cells typically do not produce GM-CSF (12, 33). Neither YFP^−^ nor YFP^+^ naïve CD4^+^ T cells produced GM-CSF after stimulation with PMA/Ionomycin ex vivo in the spleen of naïve mice, but this result was not similar in effector memory cells. However, naïve CD4^+^ T cells, both YFP^−^ and YFP^+^, did not produce IFN-γ or IL-2, which contrasted with the notable numbers of effector memory cells that produced them. This suggested that YFP^+^ naïve CD4^+^ T cells are truly naïve. Further, we sorted YFP^−^ and YFP^+^ CD4^+^ T cells and activated them with anti-CD3 and anti-CD28 mAbs for several days. Both subpopulations behaved highly similarly as regards to production of various cytokines, including GM-CSF, whereas they were notably dissimilar to effector memory cells. One possible explanation is that these YFP□ naïve CD4□ T cells are TSCM cells; however, we found no significant differences in TSCM markers (34) expression between YFP□ and YFP□ naïve CD4□ T cells. To further investigate, we compared these two cell subsets by RNA-seq. While numerous genes were differentially expressed (>900) between YFP□ and YFP□ effector/memory CD4□ T cells, YFP□ and YFP□ naïve CD4□ T cells differed in the expression of only a small set of genes. Specifically, YFP□ naïve CD4□ T cells exhibited higher expression of *Chn2* and *Crybg3* and lower expression of *Cd8b1* compared to their YFP□ counterparts. However, no well-established correlation exists between these genes and GM-CSF expression in the literature. Overall, our findings suggest that YFP□ CD4□CD62L□CD44□ T cells are genuinely naïve. The mechanisms whereby they expressed GM-CSF at some point remain unclear, but it appears that these cells did not undergo widespread and permanent changes in their phenotype. Since we observed no YFP^+^ naïve CD8^+^ T cells, this supports the view that YFP expression in some naïve CD4^+^ T cells is not an artifact of a “leaky” system. Future studies could explore whether some naïve CD4^+^ T cells, possibly those that express GM-CSF, also express other cytokines, such as IFN-γ, or if GM-CSF is unique in this regard.

Unexpectedly, we observed a subset of YFP□ Tregs in all organs analyzed. Given that GM-CSF production by Tregs has not been well characterized, we further examined YFP^+^ Tregs. A small percentage of YFP□ Tregs of naïve mice and a higher percentage of YFP□ Tregs of EAE mice expressed GM-CSF, IFN-γ, and IL-17 at levels notably higher than YFP□ Tregs. This suggests the potential plasticity of Tregs toward Th1-like or Th17-like Tregs (35, 36, 37) or the presence of pTregs (38). While pTregs are known to produce GM-CSF, IFN-γ, IL-17, and TNF (38), we found similar proportions of YFP□ and YFP□ cells expressing Helios and neuropilin-1, markers predominantly associated with tTregs rather than pTregs (27). Thus, it is more plausible that YFP□ Tregs are functionally distinct subsets of Treg cells, as Tregs could express cytokines associated with Th lineages, including IFN-γ, IL-17, and GM-CSF, particularly under inflammatory conditions (26, 37, 39). Luo et al. reported that a small percentage (1-3%) of Tregs in the spleen or spinal cord of mice with EAE express GM-CSF and IL-17 and that Tregs deficient in the transcription factor Blimp1 exhibited increased production of GM-CSF and IL-17 compared to WT Tregs (40). Interestingly, while these Tregs maintain their ability to produce some inflammatory cytokines such as IFN-γ and TNF-α in vivo, their GM-CSF expression appears to be selectively downregulated under inflammatory conditions (38). This aligns with our observation that GM-CSF expression in YFP□ Treg cells is downregulated as the disease progresses to the chronic phase. In conclusion, a greater proportion of YFP^+^ Tregs produces GM-CSF and other proinflammatory cytokines in EAE than in naïve mice. It is also notable that YFP^+^ Tregs consistently produce more of these cytokines than YFP□ Tregs, both in a steady state and EAE. In this regard, Tregs resemble YFP^+^ and YFP□ conventional effector CD4□ T cells.

Previous studies have suggested a link between CXCR6 expression and GM-CSF production in CD4□ T cells under neuroinflammatory conditions (10, 22). In this study, considerably more CD4□ YFP□ T cells were CXCR6□ than their YFP counterparts. CD4□ T cells in the liver and CNS exhibited the highest CXCR6 expression among YFP□ cells. However, during EAE, the CXCR6^+^ population increased in the spleen, lung, and CNS, suggesting a correlation between CXCR6 and GM-CSF expression, as well as T-cell trafficking and retention in inflamed tissues (20, 21, 22). Komuczki et al. reported that nearly half of CNS-infiltrating CD44□ Th cells were ex-GM-CSF□, with most co-expressing TNF, IFN-γ, and IL-17. GM-CSF□ Th cells were predominantly CXCR6 and belonged to a multi-cytokine-producing subset characterized by TNF and IFN-γ co-expression (10). Other studies indicate that CD4□CXCR6□T cells represent terminally differentiated effectors, serving as key cytokine producers in inflamed tissues. These cells proliferate rapidly and express high levels of cytolytic granule proteins (41, 42). Interestingly, nearly all CD4□ T cells that produced two or more cytokines, such as IL-17, GM-CSF, or IFN-γ, were CXCR6. Additionally, half of IL-17 single-producers and a third of GM-CSF single-producers in the LNs of EAE mice were CXCR6 (41). Consistent with previous reports, our comparison of cytokine expression between CD4□YFP□ and YFP□ cells revealed that YFP cells, both in naïve and EAE mice, exhibited a greater propensity to produce GM-CSF, IL-17, and IFN-γ and express CXCR6. We observed a strong correlation between YFP expression and the production of these cytokines in the liver, lung, and CNS of naïve mice, with their frequency further increasing during EAE. Collectively, our findings confirm that CD4□YFP□T cells constitute a multi-cytokine-producing subset characterized by high CXCR6 expression and production of GM-CSF, IFN-γ, TNF, and IL-17, with minimal IL-4 and IL-10 expression.

Although CD4^+^ effector T cells could produce GM-CSF following activation, this was not sustained in most effector cells. CD4□ T cells within the CNS of EAE mice expressed GM-CSF from the onset to the peak of the disease. However, as the disease progressed into the chronic phase, CD4□ T cells gradually lost their ability to produce GM-CSF, even upon stimulation with PMA/Ionomycin. Consistent with this, in vitro-generated ThGM cells also declined GM-CSF production after 72 hours post-activation. Komuczki et al. also reported that CNS-invading CD4□ T cells initially expressed GM-CSF, but expression declined after 14 d.p.i. (10). They reported that nearly half of CNS-infiltrating effector CD4^+^ T cells were ex-GM-CSF, while just 8% of effector CD4^+^ T cells exhibited ongoing GM-CSF expression (10). Despite this loss, the frequency of ex-GM-CSF cells remained high, indicating a gradual downregulation of GM-CSF expression. Furthermore, our follow-up analysis at 27 d.p.i., corresponding to the early chronic phase, showed further accumulation of YFP GM-CSF CD4□ T cells. These results suggest that GM-CSF expression is transient and tightly regulated, with only a subset of CD4□ T cells maintaining the ability to produce GM-CSF at any given time. This regulation likely serves as a protective mechanism to prevent excessive and potentially harmful inflammation.

In the ‘Hub and Spoke’ immune system model, T cells are licensed in the lung before circulating throughout the body (43). Increasing evidence suggests that the lung could be a key site of immune regulation in neuroinflammatory disorders (44, 45). We found that in naïve mice, CD4^+^ T cells in lymphoid organs and lungs predominantly exhibit a naïve phenotype. Following EAE induction, a shift toward an effector memory phenotype occurred across all organs, with the most pronounced changes seen in the lung and CNS. Tracking MOG --specific CD4^+^ T cells revealed a higher frequency of antigen-specific CD4□CD44□T cells in the lung compared to the spleen and liver. Odoardi et al. reported that in EAE, lung-resident effector CD4^+^ T cells expand and undergo gene-expression reprogramming, downregulating activation markers (CD25, OX40) while up-regulating molecules involved in migration (CXCR3, CCR5), adhesion (Ninj1, VLA-4), and trafficking to the CNS (44). We found that CXCR6, a marker of cytotoxic Th17 cells (22), was upregulated in lung CD4^+^ T cells following EAE, linking lung Th17 cells to neuroinflammation. In mice with EAE, CD4^+^ T cells exhibited increased IL-17 and GM-CSF expression, particularly in the lung and CNS. Notably, lung CD4^+^ T cells produced more IL-17 ex vivo following TCR activation. Our results and the literature support the view that the lung serves as a niche for both activated Th17 cells and resting myelin-reactive memory CD4^+^ T cells, which, upon stimulation, acquire migratory properties toward the CNS (44, 46).

In conclusion, this study revealed a complex and dynamic landscape of GM-CSF expression by primarily immune cells. Notably, while myeloid cells, particularly macrophages, and T cells were major GM-CSF producers, their relative contributions varied substantially between organs. The ability of individual cells to produce GM-CSF was largely transient, as the vast majority of cells that previously produced GM-CSF failed to produce it again upon activation. Surprisingly, effector memory Th cells that produced GM-CSF had a dramatically different transcriptome from those that did not. This indicates that the factors that drive GM-CSF expression also give rise to numerous other phenotypic traits. The history of GM-CSF expression in some naïve CD4^+^ T cells is a novel and intriguing observation that warrants further investigation. Furthermore, a subset of Tregs, particularly in inflammatory conditions, can acquire a Th-like phenotype characterized by the production of GM-CSF and other cytokines. Finally, we confirmed a strong association between CXCR6 expression and GM-CSF production in CD4^+^ T cells, linking this marker to a multi-cytokine-producing effector subset enriched in inflamed tissues. Future studies should focus on elucidating the molecular mechanisms governing the transient nature of GM-CSF expression, the origin and relevance of GM-CSF expression in naïve CD4^+^ T cells, and the potential role of GM-CSF production by Tregs.

## Material and methods

### Sex as a biological variable

Although sex was not considered as a biological variable, approximately equal numbers of male and female mice aged 2-3 months were used in the experiments, which were conducted with prior approval from the Institutional Animal Care and Use Committee of Thomas Jefferson University.

### Mice

The GM-CSF reporter (*Gr*) transgenic mouse line was generated in-house. *Gr* mice carry a transgenic GM-CSF (*Csf2*) allele that enables normal GM-CSF expression, along with iCre and blue fluorescent protein (BFP) expression (29). To create a GM-CSF reporter/fate reporter (*Gr/fr*) double transgenic line, we crossed *Gr* mice with Rosa26eYFP mice (The Jackson Laboratory). *Gr/fr* mice enable lineage tracing through iCre-induced continuous yellow fluorescent protein (YFP) expression. Thus, cells with ongoing GM-CSF expression are BFP^+^ (reporting function), while cells that previously expressed GM-CSF are permanently YFP^+^ (fate reporting function) (8, 29). Since we stained for multiple cytokines in parallel, we opted to stain cells with anti-GM-CSF monoclonal antibody (mAb) instead of solely relying on BFP expression to identify GM-CSF-expressing cells. Nonetheless, in some experiments, we relied on BFP expressions to identify GM-CSF^+^ cells, and **Fig. S1A** demonstrates an excellent correlation between BFP and GM-CSF expression.

In our preliminary studies, we noticed that *Gr/fr* mice with one *Csf2*/iCre/BFP allele and one wild type (WT) *Csf2* allele had a lower range (6-13%; mean 8%) of YFP^+^CD4^+^ T cells in peripheral blood compared to mice with two *Csf2*/iCre/BFP alleles (7-22%; mean 13%). This indicated that not all cells with one *Csf2*/iCre/BFP allele and one WT *Csf2* allele became YFP^+^ upon GM-CSF expression. Even though GM-CSF expression is not considered to be strictly monoallelic, various factors can lead to differential expression of alleles within the same cell (47). Hence, heterozygous transgenic cells that express GM-CSF primarily from the WT *Csf2* allele would fail to become YFP^+^ because they would not express enough iCre from the *Csf2*/iCre/BFP allele to activate YFP expression. Therefore, to ensure that the maximum percentage of cells expressing GM-CSF become YFP^+^, we used *Gr/fr* mice that were homozygous for both the transgenic *Csf2*/iCre/BFP allele and the Rosa26eYFP allele.

Naïve *Gr/fr* mice, aged 6 weeks, exhibited YFP expression in 13 ± 8 % of CD4^+^ T cells from peripheral blood. We included all the mice in experiments that analyzed YFP expression in various cell types from different organs, as described in the first section of the results. In experiments focused on the CD4^+^ and CD8^+^ T cell subsets and functional analysis, we used mice with 10-15% YFP^+^CD4^+^ T cells to ensure a more homogeneous sample.

*Cxcr3□*/□and *Cxcr6□*/□mouse lines with the C57BL/6 background and WT C57BL/6 (B6) mice were purchased from the Jackson Laboratory.

### Mononuclear cell isolation

Mice were anesthetized using a ketamine-xylazine cocktail. After blood collection, mice were perfused through the left ventricle with cold PBS. Then, inguinal LNs, spleen, BM, and thymus were collected and mechanically dissociated. The central nervous system (CNS), liver, small intestine (SI), kidney, and lung were collected and digested with DNase (50 µg/ml; Stem Cell) and Liberase (500 µg/mL for CNS; 200 µg/mL for liver, kidney, and SI; 100 µg/ml for lung; Sigma-Aldrich), in RPMI for 30 min at 37 °C, followed by mechanical dissociation. The debris removal solution (Miltenyi Biotec) efficiently removed cell debris from the liver, kidney, and lungs. For the CNS, myelin was removed using a 40% Percoll (Cytiva) gradient at 2000 RPM for 30 min at 24 °C. SI cells were suspended in a 40% Percoll (Cytiva) overlaid on a 70% Percoll and spun at 2000 RPM for 25 min at 24 °C. Mononuclear cells (MNCs) were collected from the interface between Percoll layers. Peripheral blood mononuclear cells (PBMCs) were isolated from whole blood by Ficoll density gradient centrifugation.

### Flow cytometry and intracellular staining

Cells were activated with PMA (50 ng/ml; Sigma-Aldrich), Ionomycin (500 ng/ml; Sigma-Aldrich), and GolgiPlug (1 µg/ml; BD Biosciences) at 37 °C for 4-6 h. Cells were washed, and anti-CD16/32 Ab (eBioscience) was used to block the non-specific binding of Abs to Fc receptors on myeloid cells. The cells were then stained with Abs for surface markers (**Table S1**). After fixation/permeabilization with Caltag Fix/Perm reagents (Invitrogen), cells were stained with Abs to detect intracellular markers (**Table S1**). Data was acquired on FACSAria Fusion II (BD Biosciences) and analyzed by FlowJo software (TreeStar).

### MHC Tetramer Staining

MNCs were isolated from the spleen, liver, and lung of MOG_35-55_-immunized mice at 8 d.p.i., followed by CD4^+^ T cell enrichment using MACS. A total of 5 × 10 cells were first stained with I-Ab MOG_35-55_ MHC tetramer-PE (MBL, Japan), CD3, CD4, and CD44 and then analyzed by flow cytometry.

### CD4^+^ T cells isolation, co-culture, and Th differentiation

Various subpopulations of CD4^+^ T cells from multiple organs of 2-3-month-old *Gr/fr* mice were sorted via fluorescence-activated cell sorting (FACS) using a FACSAria fusion II flow cytometer system (BD Biosciences). Flow cytometry confirmed the purity of sorted cells. T cell-depleted splenocytes were prepared using anti-CD3 beads and magnetic-activated cell sorting (MACS). The sorted CD4^+^ T cells were co-cultured with the T cell-depleted splenocytes at a 1:4 ratio in the presence of soluble anti-CD3 and anti-CD28 mAbs (3 µg/mL) for 3 days. After co-culture, supernatants were collected for cytokine measurement using enzyme-linked immunosorbent assay (ELISA), while cells were stimulated with PMA, Ionomycin, and GolgiPlug for 5 h and analyzed by flow cytometry.

For Th differentiation, naïve (CD62L^hi^CD44^−^) YFP□ CD25^−^CD4^+^ T cells from the spleens of *Gr/fr* mice were FACS sorted and differentiated into ThGM and Th1 cells, as previously described (8). Briefly, naïve T cells were cultured at a ratio of 1:4 with T cell-depleted splenocytes at a density of 1×10^6^ cells/ml. Naïve T cells were activated with soluble anti-CD3 and anti-CD28 mAbs (3 µg/mL) for 3 days in different Th differentiation conditions: ThGM: anti-IFN-γ (10 μg/ml), anti-IL-12 (10 μg/ml), anti-IL-4 (5 μg/ml) Abs, and Th1: IL-12 (20 ng/ml) Ab.

### Bulk mRNA sequencing

YFP^+^, YFP^−^ naïve (CD62L^+^CD44^−^), and T_EM_ (CD62L^−^CD44^hi^) CD4^+^ T cells from the spleens of *Gr/fr* mice (n=3) were FACS sorted. Then, RNA was extracted using RNeasy Mini Kit (Qiagen) according to the manufacturer’s instructions. Libraries were prepared using 200 ng of total RNA and the Illumina TruSeq Stranded Total RNA library preparation kit. Libraries’ qualities were evaluated using the PerkinElmer Labchip GX and qPCR using the Kapa Library Quantification Kit and the Life Technologies Viia7 Real-time PCR instrument. Libraries at a concentration of 2nM were sequenced on the Illumina HiSeq2500 using the High Output v4 chemistry. Raw FASTQ sequencing reads were mapped against the mm10 reference genome using the DRAGEN genome pipeline. BAM files were used to generate a feature/gene counts matrix using the R/Bioconductor package Rsubread. Differential expression contrasts were performed using the DESeq2 package in R/Bioconductor. Differentially expressed genes (DEGs) were identified as those with Benjamini-Hochberg adjusted p ≤ 0.05 and absolute log2 fold change ≥ 2. All plots were generated using R/Bioconductor.

### Statistical analysis

Statistical analysis was conducted using GraphPad Prism 8 software. Data was analyzed using an unpaired, two-tailed Student’s t-test between two groups and a one-way ANOVA between three or more groups. P ≤ 0.05 was considered significant. Data represent mean ± SEM.

## Availability of data and materials

The datasets used and/or analyzed in the current study are available from the corresponding author upon request.

## Acknowledgments

We thank Hemma Murali, Scholarly Writing Specialist at Thomas Jefferson University.

## Author information

### Authors and Affiliations

Gholamreza Azizi, Javad Rasouli, Hamed Naziri, Michael V. Gonzalez, James Garifallou, Guang-Xian Zhang, Bogoljub Ciric, Abdolmohamad Rostami

Department of Neurology, Thomas Jefferson University, Philadelphia, PA, 19107, USA.

The Center for Applied Genomics, The Children’s Hospital of Philadelphia, Philadelphia, PA, USA.

Division of Neonatology, Children’s Hospital of Philadelphia, Philadelphia, PA, USA.

### Contributions

BC, GXZ, and AR conceived the study and obtained funding for it. GA, JR, HN, MVG, and JG performed experiments. All authors contributed to the study plan, interpreted data, and wrote and reviewed the manuscript.

### Corresponding author

Correspondence to Abdolmohamad Rostami

## Ethics declarations

### Ethics approval and consent to participate

All animal experiments were performed with the approval of the Institutional Animal Care and Use Committee (IACUC) at Thomas Jefferson University.

## Supplemental material

**Figure S1.**
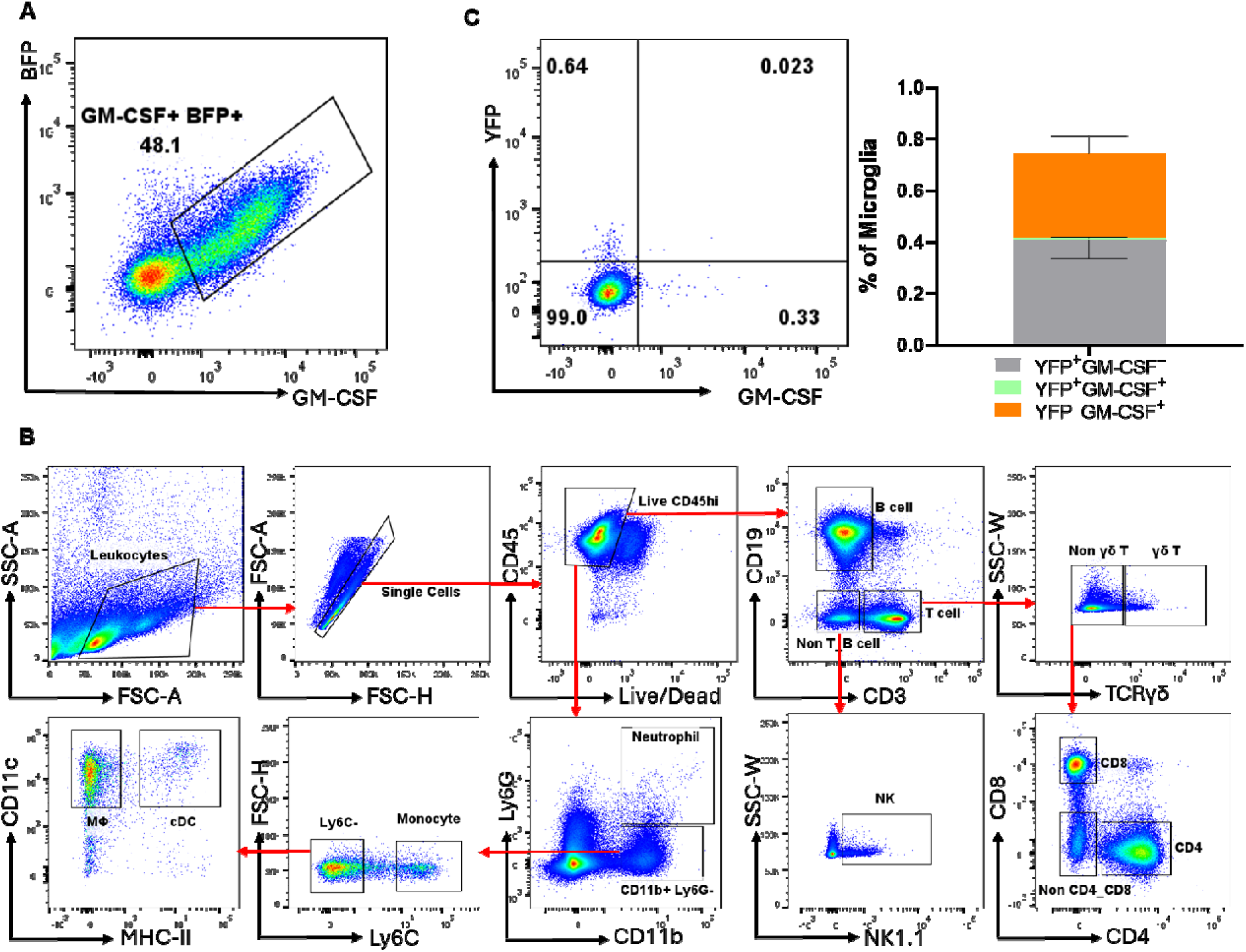
Flow cytometric analysis and gating strategy for immune cell subsets and GM-CSF expression in *Gr/fr* Mice. **(A)** Correlation between BFP and GM-CSF expression in CD4^+^ T cells isolated from the CNS of mice with EAE. Mononuclear cells were obtained from the CNS of *Gr/fr* mice at the peak of EAE. The cells were stimulated for 6 h with PMA, Ionomycin, and GolgiPlug, followed by staining for surface and intracellular antigens. **(B)** Gating strategy for flow cytometric analysis of lymphoid and myeloid cells among splenocytes of *Gr/fr* mice. A similar gating strategy was used in analyses of immune cells from other organs. **(C)** Representative flow cytometry dot plot and a stacked bar chart show YFP and GM-CSF expression by gated microglial cells from the CNS of mice with EAE. MΦ: macrophage, cDC: conventional dendritic cell.

**Figure S2.**
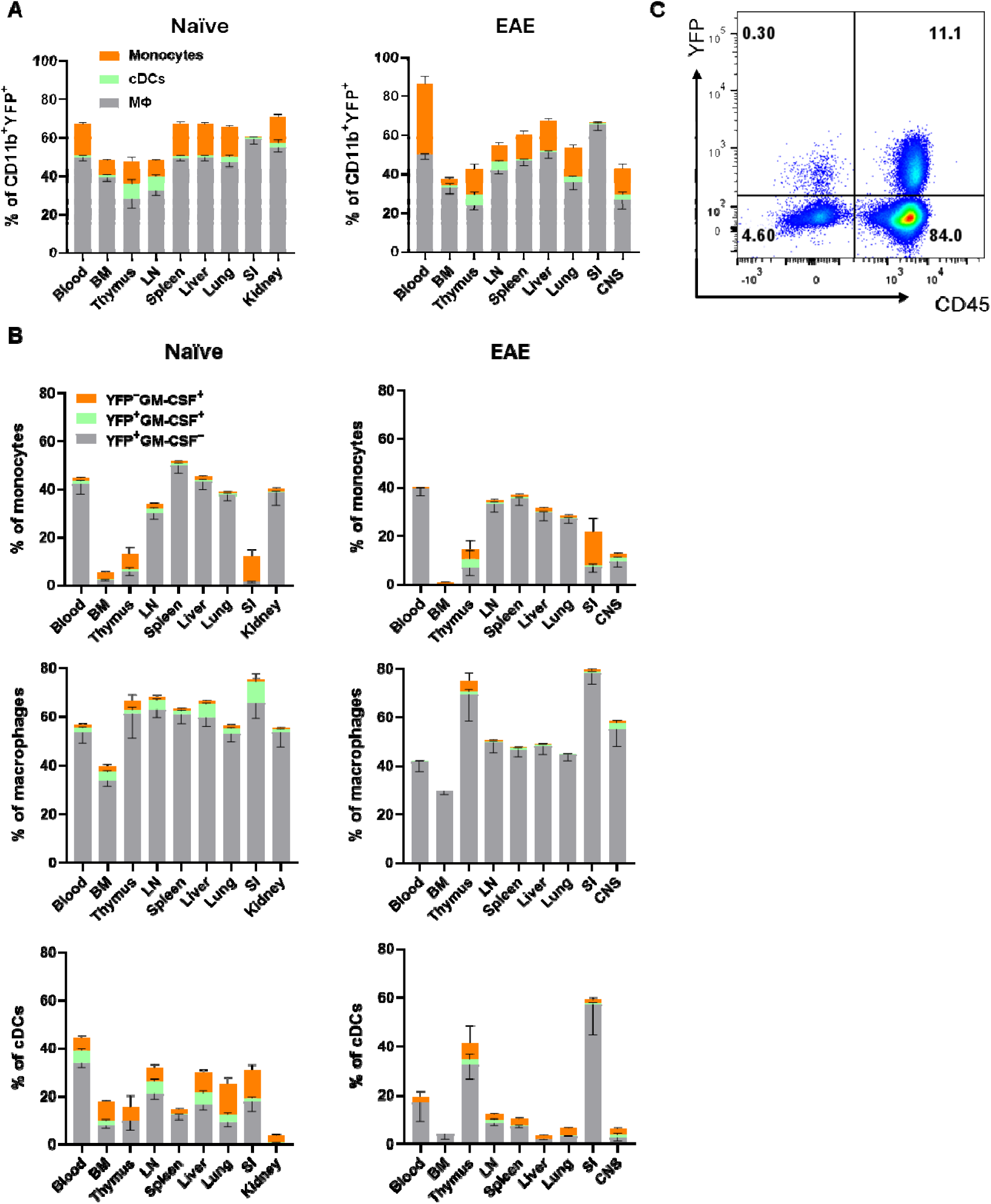
GM-CSF and YFP expression in CD11b^+^ subsets from mouse organs. Mononuclear cells were isolated from the blood, bone marrow (BM), thymus, lymph nodes (LNs), spleen, liver, lung, small intestine (SI), and kidney of naïve mice, as well as from the CNS of mice with EAE at peak clinical disease severity. Mice used were 2-3 months old, male and female *Gr/fr*. Cells were stimulated with PMA, Ionomycin, and GolgiPlug, stained for CD45, myeloid-specific surface markers, and GM-CSF, and analyzed by flow cytometry. YFP and GM-CSF expressions were quantified within gated live CD45^hi^ populations. **(A)** Proportions of major myeloid cell types among CD45^hi^CD11b^+^YFP+ immune cells in naïve and EAE mice. Frequencies of YFP^+^ cells were determined within gated live CD45^hi^CD11b^+^ populations. Stacked bar charts display the proportions of major myeloid cell types among these YFP^+^ cells. **(B)** The frequencies of YFP^+^ and GM-CSF^+^ cells among major myeloid cell types in naïve mice and mice with EAE. **(C)** An example of YFP expression in CD45^+^ and CD45^-^ cells among live cells (from the lung of EAE mice). Data are presented as the mean ± SEM from two independent experiments in naïve (total n = 13) and EAE (total n = 8) mice. cDC: conventional dendritic cells, MΦ: macrophages. Abbreviations: LN- lymph nodes, BM- bone marrow, SI- small intestine, CNS- central nervous system.

**Figure S3.**
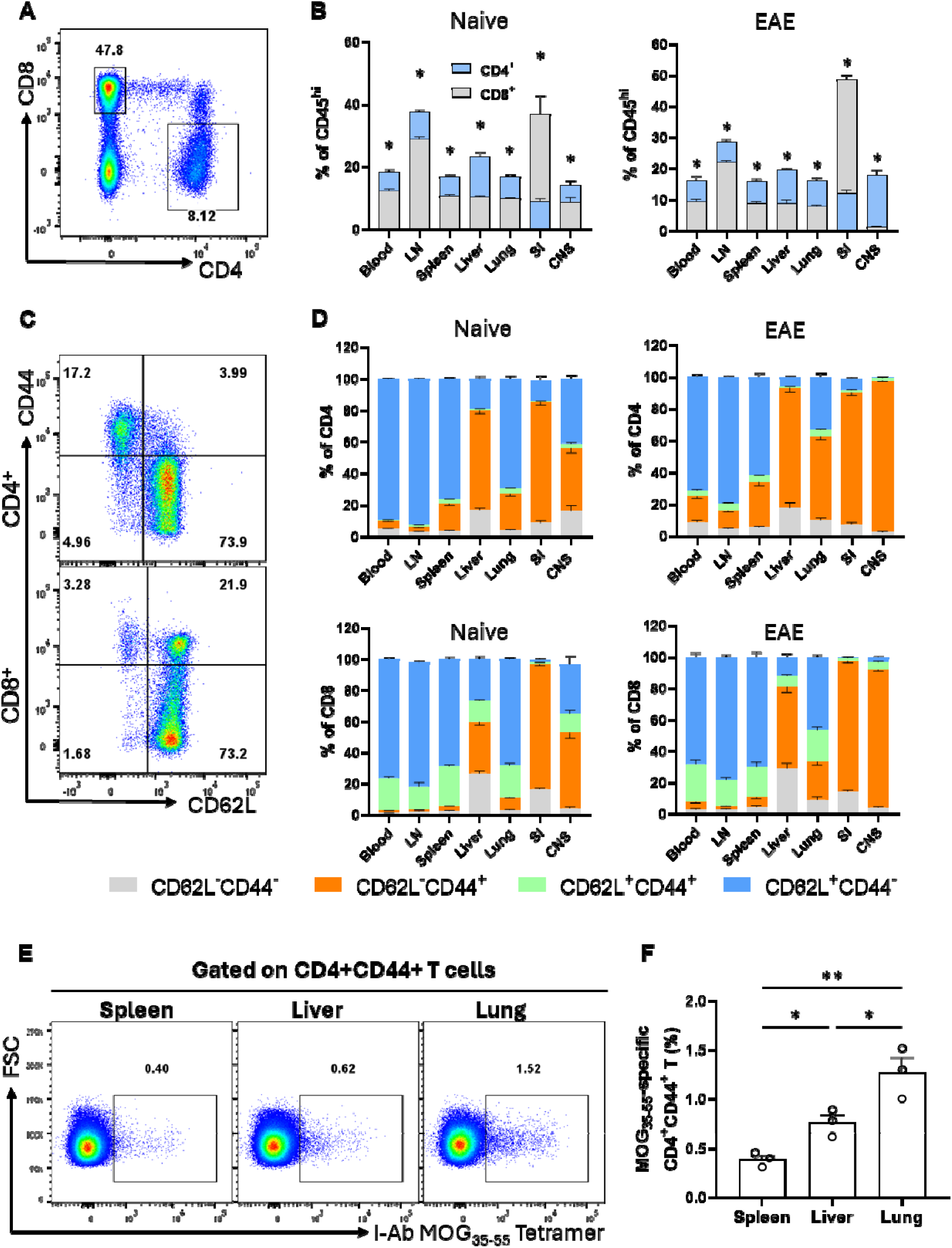
CD4^+^ and CD8^+^ T cell subset frequencies in mouse organs. Both naïve and EAE mice (clinical score 2-3) male and female *Gr/fr* mice aged 2-3 months were sacrificed. Their cells were isolated from the blood, LNs, spleen, liver, lung, SI, and CNS, stained and analyzed by flow cytometry for CD45, CD4, CD8, CD62L, and CD44 expression. **(A)** An example of the gating strategy for CD4^+^CD8^−^ and CD4^−^CD8^+^ cells within live CD45^hi^ cells (from the SI of naïve mice). **(B)** Superimposed bar charts illustrate CD4^+^ and CD8^+^ T cell frequencies across multiple organs of naïve and EAE mice. **(C)** Examples of gating strategy for CD62L^−^CD44^−^, CD62L^+^CD44^−^, CD62L^−^CD44^+^, and CD62L^+^CD44^+^ subsets within conventional (Foxp3^−^) CD4^+^ T cells (upper panel) and CD8^+^ T cells (lower panel) from the spleen of naïve mice. **(D)** Stacked bar charts show frequencies of CD62L^−^CD44^−^, CD62L^+^CD44^−^, CD62L^−^CD44^+^, and CD62L^+^CD44^+^ subsets within total conventional (Foxp3^−^) CD4^+^ and CD8^+^ T cell populations from naïve and EAE mice. **(E)** Cells from the spleen, liver, and lung of MOG□□□□□-immunized mice were harvested at 8 dpi, stained with I-A□ MOG□□□□□ MHC tetramer for CD3, CD4, and CD44 markers before analysis by flow cytometry. Representative dot plots for the spleen, liver, and lung illustrate the percentage of tetramer□ cells within the CD4□CD44□ T cell population. **(F)** The frequencies of MOG□□□□□-specific CD4□CD44□ T cells among total CD4□CD44□ T cells are presented as mean ± SEM (n = 3 per group per experiment), based on data from two independent experiments. Data represent two independent experiments with naïve (total n = 7) and EAE (total n = 8) mice for panels A–D. Statistical analysis was performed using Student’s t-test; p < 0.05. Abbreviations: LN- lymph nodes, SI- small intestine, CNS- central nervous system.

**Figure S4.**
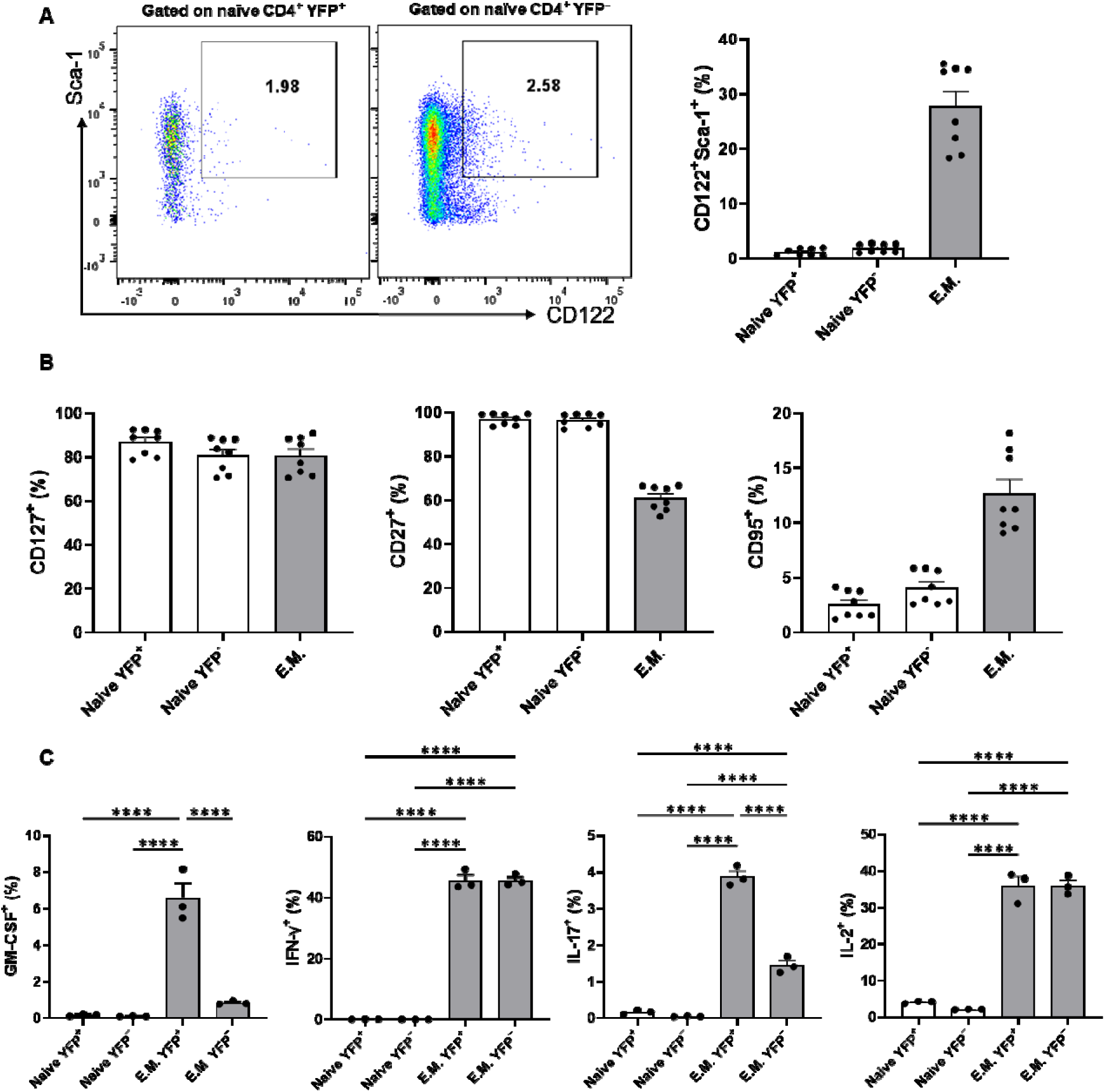
Comparison of naïve (CD62L□CD44^−^) YFP□ and YFP^−^ CD4□ T cells. CD4□ T cells were isolated from the spleens of naïve 2- to 3-month-old male and female *Gr/fr* mice using magnetic-activated cell sortin (MACS). Cells were stained for CD4, CD62L, CD44, CD122, Sca-1, CD95, CD27, and CD127, and analyzed b flow cytometry. **(A)** The gating strategy for Sca-1LCD122L cells in gated live naïve (CD62L□CD44^-^) YFP□ (left) and YFP^−^ (right) CD4□ T cells, along with their corresponding bar graph with summarized data from multiple mice. **(B)** Expression of CD127, CD27, and CD95 in naïve (CD62L□CD44^−^) YFP□ and YFP^−^ CD4□ T cells. Effector memory (CD62L^−^CD44^+^) CD4□ T cells were included for comparison. (**C**) The cytokine expression b YFP□ and YFP− populations of naïve and effector memory CD4□ T cells in the spleens of naïve 2-3-month-old female *Gr/fr* mice. Cells were stimulated with PMA, Ionomycin, and GolgiPlug for 5 h, followed by staining for surface and intracellular antigens. Data are presented as mean ± SEM from two independent experiments for A an B (total n = 8) and a single experiment for C (n=3). Statistical analysis was conducted using an unpaired Student’s t-test and one-way ANOVA. ****P < 0.0001.

**Figure S5.**
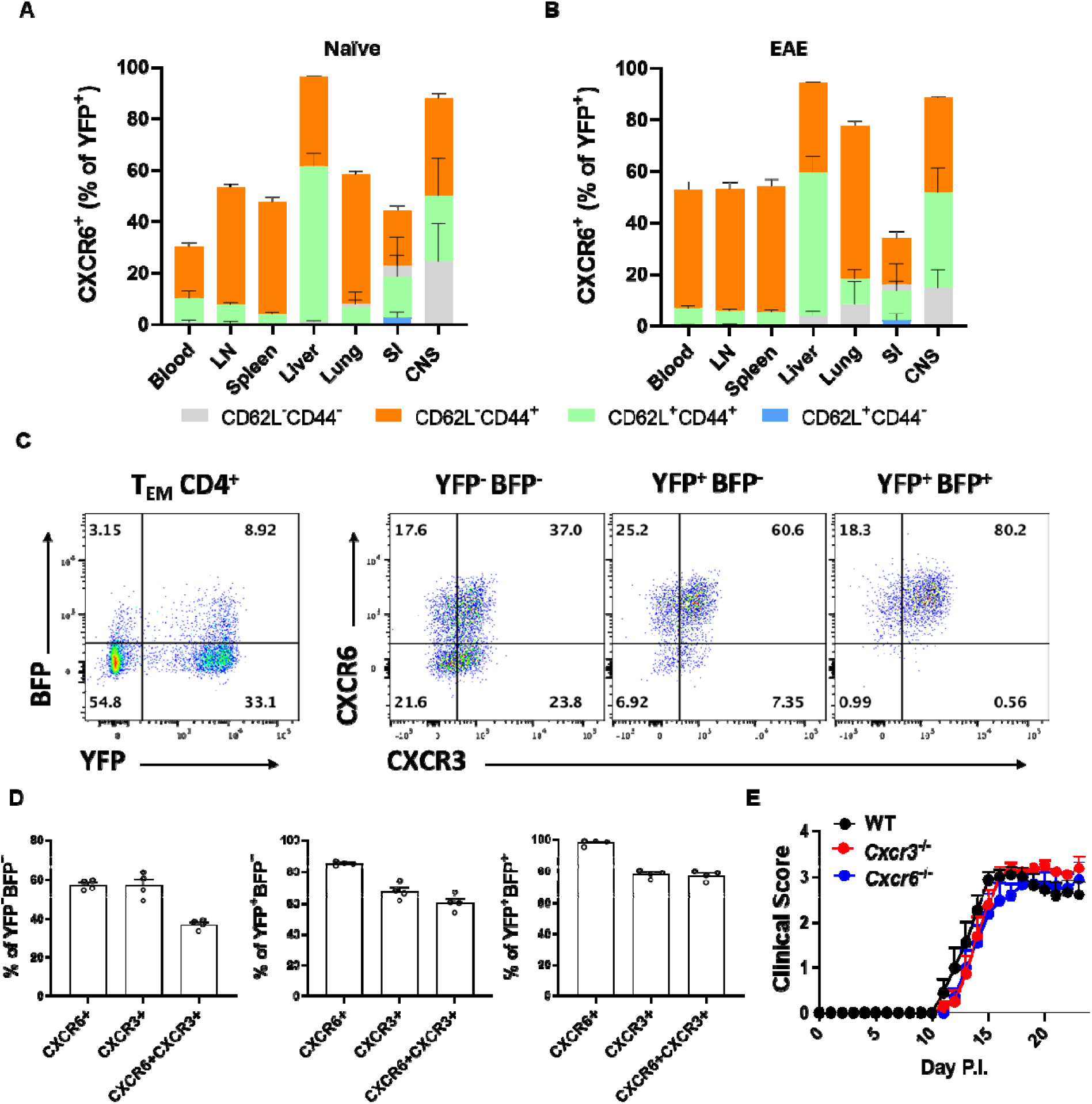
CXCR6 and YFP expression in CD4^+^ T cell subsets. Mononuclear cells were isolated from the blood, LNs, spleen, liver, lung, SI, and CNS of 2–3-month-old male and female *Gr/fr* mice. Cells were stained for CD45, CD4, CD62L, CD44, and CXCR6. Conventional (Foxp3^−^) CD4□ T cell subsets were defined based on CD62L and CD44 expression, and YFP□ cells were analyzed for CXCR6 expression in **(A)** naïve and **(B**) EAE mice, shown in superimposed bar charts. **(C, D)** Mononuclear cells were isolated from the CNS of mice with EAE (15 dpi) and stained for surface markers. Flow cytometry plots show YFP, BFP, CXCR6, and CXCR3 expression within T_EM_ (CD62L^−^CD44^hi^) CD4□ cells. **(E)** Clinical scores of *Cxcr6*□/□, *Cxcr3*□/□, and WT mice immunized for EAE induction. The data represents two independent experiments, with naïve (total n = 7) and EAE-induced (total n = for panel B and total n=10 for panel E) mice and n = 4 from one experiment for C, and D. Error bars indicate mean ± SEM.

**Figure S6.**
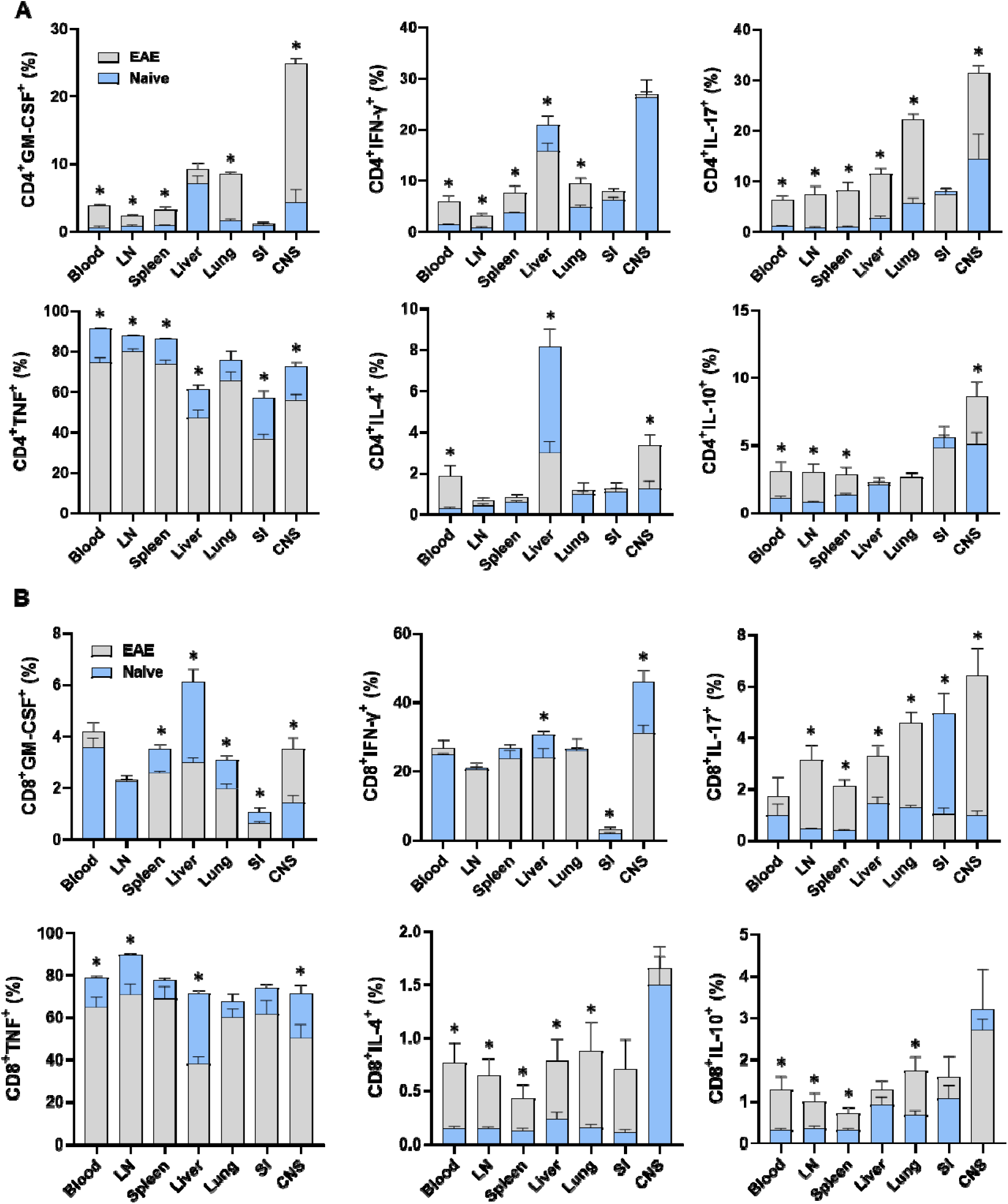
The cytokine expression by CD4^+^ and CD8^+^ T cells from different organs. Mononuclear cells wer isolated from the blood, LN, spleen, liver, lung, SI, and CNS of 2–3-month-old male and female *Gr/fr* mice, naïve and with EAE. Cells were stimulated with PMA, Ionomycin, and GolgiPlug, and stained for the surface an intracellular antigens of **(A)** CD4^+^ and **(B)** CD8^+^ T cells, shown in superimposed bar charts. Results are expressed as the mean ± SEM with naïve (total n = 7) and EAE-immunized (total n = 8) from 2 independent experiments. Statistical analysis was performed using Student’s t-test; *P<0.05.

**Figure S7.**
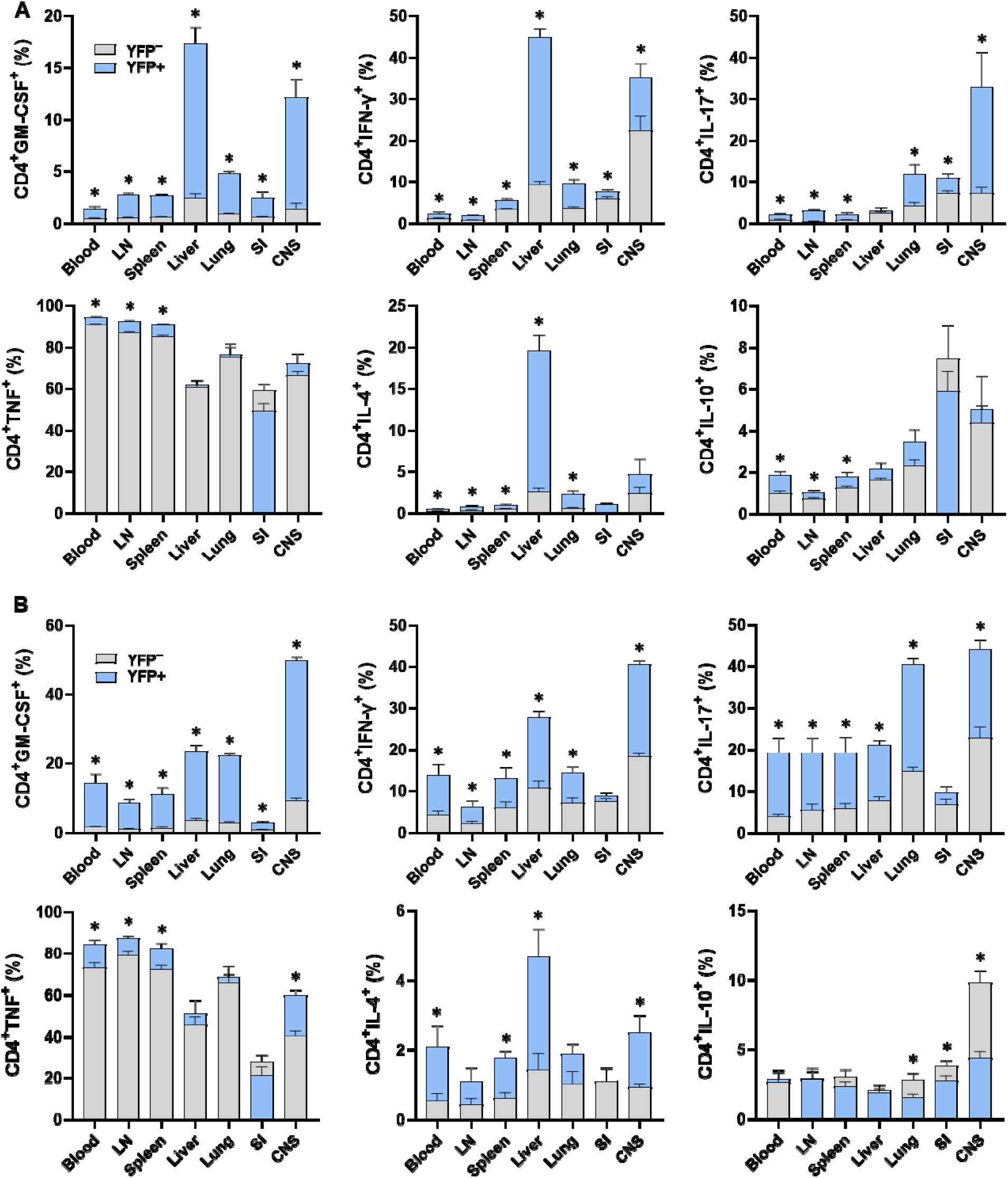
The cytokine expression by YFP^+^ and YFP^−^ CD4^+^ T cells. Mononuclear cells were isolated from the blood, LNs, spleen, liver, lung, SI, and CNS of 2–3-month-old male and female naïve and EAE *Gr/fr* mice. Cells were stimulated with PMA, Ionomycin, and GolgiPlug and stained for surface and intracellular antigens. Superimposed bar charts illustrate GM-CSF, IFN-γ, IL-17, TNF, IL-4, and IL-10 expression by gated YFP^+^ and YFP^−^ CD4^+^ T cells from **(A)** naïve and **(B)** EAE mice. Results are expressed as the mean ± SEM; naïve (total n = 7) and EAE-immunized (total n = 8) from 2 independent experiments. Statistical analysis was performed using Student’s t-test; *P<0.05.

**Figure S8.**
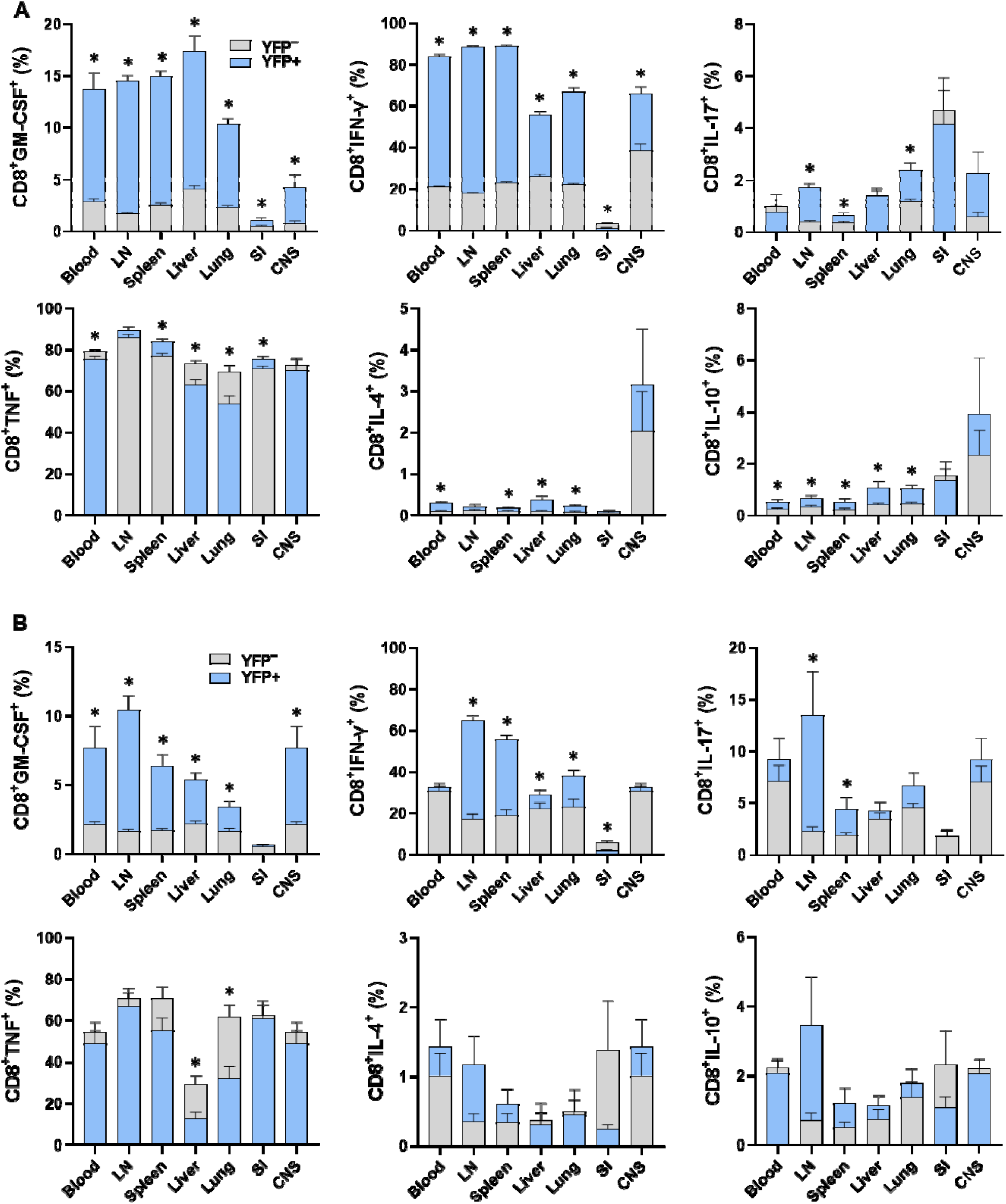
The cytokine expression by YFP^+^ and YFP^−^ CD8^+^ T cells. Mononuclear cells were isolated from the blood, LNs, spleen, liver, lung, SI, and CNS of 2-3-month-old male and female naïve and EAE *Gr/fr* mice. Cells were stimulated with PMA, Ionomycin, and GolgiPlug and stained for surface and intracellular antigens. GM-CSF, IFN-γ, IL-17, TNF, IL-4, and IL-10 expression by gated YFP^+^ and YFP^−^ CD8^+^ T cells from **(A)** naïve and **(B)** EAE mice is shown using superimposed bar charts. Results are expressed as the mean ± SEM; naïve mice (total n = 7) and mice with EAE (total n = 8) from 2 independent experiments. Statistical analysis was performed using Student’s t-test; *P<0.05.

**Figure S9.**
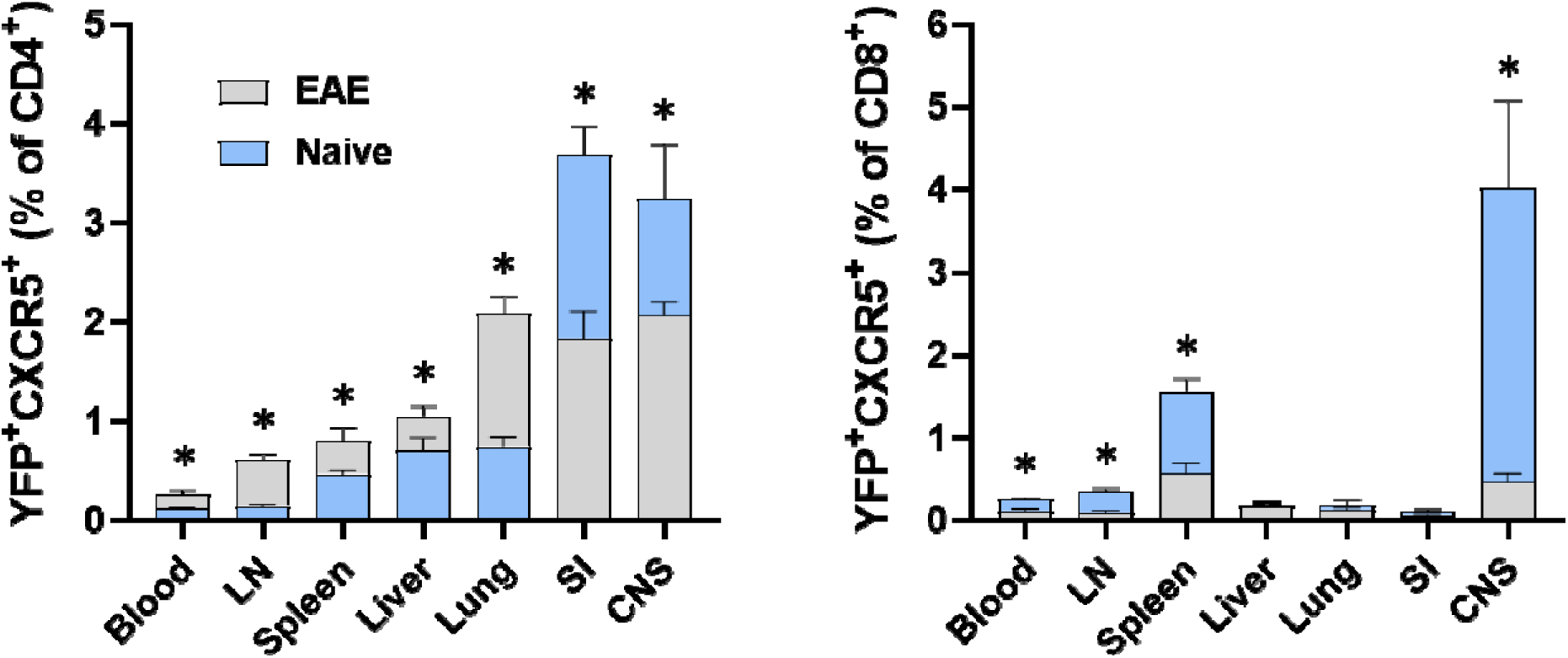
YFP and CXCR5 expression by CD4^+^ and CD8^+^ T cells. Superimposed bar charts illustrate YFP and CXCR5 expression by gated CD4^+^ or CD8^+^ T cells. Results are expressed as the mean ± SEM; naïve (total n = 7) and EAE-immunized (total n = 8) from 2 independent experiments. Statistical analysis was performed usin Student’s t-test; *P<0.05.

**Figure S10.**
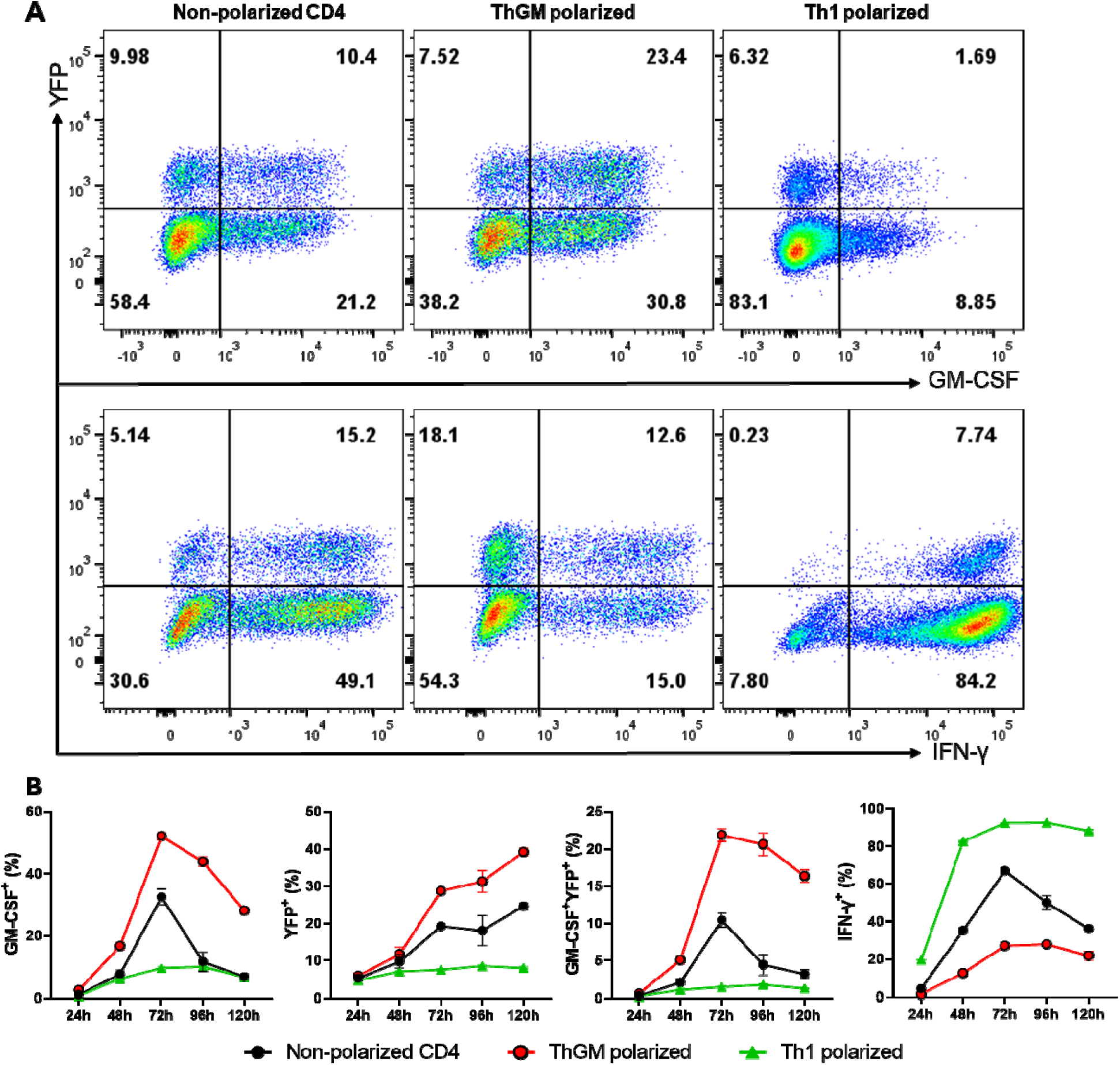
GM-CSF and YFP expression during Th cell polarization. Naïve YFP^−^CD4□ T cells (CD4□CD25^−^CD62L□CD44^lo^YFP^−^) were sorted from the spleens of naïve 2- to 3-month-old male and female *Gr/fr* mice and co-cultured with T cell-depleted splenocytes at a ratio of 1:4 in the presence of soluble anti-CD3/CD28 mAbs (3 µg/ml) under ThGM and Th1 differentiation conditions; ThGM: anti-IFN-γ (10 μg/ml), anti-IL-12 (10 μg/ml), anti-IL-4 (5 μg/ml) antibodies, and Th1: IL-12 (20 ng/ml). Cells were harvested every 24 h over a 5-day period, then stimulated with PMA, Ionomycin, and GolgiPlug, followed by staining for CD4, GM-CSF, and IFN-γ. **(A)** Flow cytometry plots illustrate gating for GM-CSF, IFN-γ, and YFP expression in gated CD4□ T cells after 72h stimulation. **(B)** GM-CSF, IFN-γ, and YFP expression in gated CD4□ T cells derived from naïve CD4□YFP^−^ T cells over time. Data is presented as mean ± SEM from a single experiment (n=3).

**Table S1.** Anti-mouse flow cytometry antibodies.

| **Antigen** | **Fluorochrome** | **Clone** | **Supplier** | **Category** |
| --- | --- | --- | --- | --- |
| CD3 | APC/Cy7 | 17A2 | Biologend | Surface |
| CD3 | AF700 | 17A2 | Biologend | Surface |
| CD4 | AF700 | GK1.5 | Biologend | Surface |
| CD4 | BV711 | RM4-5 | Biologend | Surface |
| CD4 | BV650 | GK1.5 | Biologend | Surface |
| CD8 | PE/Cy7 | 53-6.7 | Biologend | Surface |
| CD8 | BV711 | 53-6.7 | Biologend | Surface |
| CD8 | APC/Cy7 | 53-6.7 | Biologend | Surface |
| CD19 | PE | 6D5 | Biologend | Surface |
| CD25 | BV711 | PC61 | Biologend | Surface |
| CD25 | AF700 | PC61 | Biologend | Surface |
| CD27 | BV685 | LG.3A10 | Biologend | Surface |
| CD44 | BV650 | IM7 | Biologend | Surface |
| CD44 | PE/Cy7 | IM7 | Biologend | Surface |
| CD45 | AF700 | 30-F11 | Biologend | Surface |
| CD45 | BV650 | 30-F11 | Biologend | Surface |
| CD62L | APC | MEL-14 | Biologend | Surface |
| CD62L | PE/Cy5 | MEL-14 | Biologend | Surface |
| CD95 | BV605 | SA367H8 | Biologend | Surface |
| CD122 | PE | 5H4 | Biologend | Surface |
| CD127 | APC/Cy7 | A7R34 | Biologend | Surface |
| CD304 | PerCP-eFluor 710 | 3DS304M | Invitrogen | Surface |
| Sca-1 | BV711 | D7 | Biologend | Surface |
| TCR $\gamma\delta$ | APC | QA20A46 | Biologend | Surface |
| NK-1.1 | BV785 | PK136 | Biologend | Surface |
| CXCR5 | BV785 | L138D7 | Biologend | Surface |
| CXCR6 | PE/Dazzle™ 594 | SA051D1 | Biologend | Surface |
| CD11b | BV785 | M1/70 | Biologend | Surface |
| CD11c | PE | N418 | Biologend | Surface |
| Ly6C | BV650 | HK1.4 | Biologend | Surface |
| Ly6G | BV711 | 1A8 | Biologend | Surface |
| F4/80 | PE/Cy7 | BM8 | Biologend | Surface |
| MHC-II | APC/Cy7 | M5/114.15.2 | Biologend | Surface |
| IFN-γ | PE/Cy7 | XMG1.2 | Biologend | Intercellular |
| TNF | APC/Cy7 | MP6-XT22 | Biologend | Intercellular |
| GM-CSF | PE/Dazzle™ 594 | MP1-22E9 | Biologend | Intercellular |
| IL-2 | PE | JES6-5H4 | BD Biosciences | Intercellular |
| IL-4 | APC | 11B11 | Biologend | Intercellular |
| IL-10 | PE | JES5-16E3 | Biologend | Intercellular |
| IL-17 | BV785 | TC11-18H10.1 | Biologend | Intercellular |
| Foxp3 | eFluor 450 | FJK-16S | eBioscience | Intercellular |
| Foxp3 | PE | FJK-16S | eBioscience | Intercellular |
| Helios | APC | 22F6 | eBioscience | Intercellular |

IL; Interleukin, IFN; Interferon, TNF; Tumor necrosis factor, GM-CSF; Granulocyte-macrophage colony-stimulating factor, Foxp3; Forkhead box P3, Sca-1; Stem cell antigen-1, CXCR; CXC chemokine receptor, MHC class II; Major Histocompatibility Complex class II.

